# Obligate multicellularity circumvents population genetic barriers to collective-level adaptation

**DOI:** 10.64898/2026.05.14.725289

**Authors:** Autumn Peterson, Eric Libby, Anthony Burnetti, Jenasis Campbell, William Ratcliff

## Abstract

’Complex’ multicellularity has evolved in five eukaryotic lineages, all of which develop clonally and are obligately multicellular (*i.e.*, lack a reproductive unicellular phase). Clonal development is well-established as a critical factor promoting multicellular adaptation by limiting within- group conflict, but little is known about the evolutionary consequences of obligate multicellularity itself. Here, we show that obligate multicellularity alleviates an establishment barrier to collective-level adaptation. We engineered a single strain of snowflake yeast (*Saccharomyces cerevisiae*) capable of exhibiting obligately multicellular, facultatively multicellular, or obligately unicellular life cycles, and evolved them under selection for larger group size. All five obligately multicellular populations rapidly evolved larger groups, primarily through whole genome duplication. Facultative populations were far more constrained, with tetraploidy evolving in only 2/10 populations, despite this trait being strongly beneficial across the full life cycle. Mathematical modeling suggests that group formation and opposing selection across life cycle phases interact, increasing the chance that mutations beneficial across the full life cycle are lost before they establish. These results suggest that clonality and obligate multicellularity jointly create the conditions for complex multicellularity to evolve, one by suppressing within-group conflict, the other by removing population genetic barriers to collective-level adaptation.

## Introduction

Multicellularity has evolved 45 times in eukaryotes alone^1^, generating diverse morphologies ranging from small, simple groups to large organisms consisting of many cell types^2–4^. A key step in this transition is a shift in Darwinian individuality, where selection acts primarily on the traits of groups rather than individual cells^5–9^. For this to occur, groups must possess heritable variation in multicellular traits that affect fitness, allowing open-ended adaptation in collective-level traits to occur over generations^7,10^.

The structure of a nascent multicellular life cycle can either facilitate^11–13^ or constrain^14,15^ a lineage’s capacity for multicellular adaptation^16,17^. In contrast to the many origins of simple multicellularity, ‘complex’ multicellularity has evolved in just five lineages: animals, plants, fungi, red algae, and brown algae^18,19^. These lineages share two key life cycle traits: they develop clonally and are obligately multicellular^11,20–22^. Indeed, the importance of clonality has long been understood through the lens of social evolution: within-group genetic diversity creates the potential for intra-organismal selection, which may undermine multicellular innovations^5,23–25^. Clonality also increases the heritability of multicellular traits: in clonal organisms, the group has a single underlying genome, allowing selection acting on group-level traits to simultaneously act on the genome responsible for generating them^26^. This allows even emergent multicellular traits, arising from mutations that only directly affect cellular phenotype, to be highly heritable^27^. In contrast, chimeric organisms face a fundamental disconnect between the target of selection, the multicellular phenotype, and the units of heredity, individual genomes.

Most prior work on nascent multicellular life cycles has focused on the distinction between ’staying together’ and ’coming together’^16,20,21,28–31^, which tends to conflate within-group genetic diversity arising from aggregation with facultative multicellularity. By comparison, relatively little work has directly considered the role of obligate vs. facultative multicellularity in the evolution of organismal complexity. We adopt the definition of Marquez-Zacarias et al.^24^: obligately multicellular organisms lack a persistent, reproductive unicellular phase, while facultatively multicellular organisms alternate between unicellular and multicellular phases, with growth and reproduction occurring in both states. To our knowledge, all lineages which facultatively form clonal multicellular groups have remained simple^32–35^, despite avoiding disruption by cheats. This suggests that obligate multicellularity may itself play a role in the evolution of increasingly complex organismality.

There may be profound population-genetic consequences of this life cycle trait. Few behaviors can reduce population size as dramatically as the evolution of large, clonal multicellular organisms^36–38^. For example, the average human has 37 trillion cells^39^, but counts as a single unit of selection; if those cells were instead growing autonomously, they would contribute an *N* of 37 trillion to a population. A lineage alternating between unicellular and multicellular phases changes both the fitness effects of a mutation and the number of independently sampled units. A mutation that increases group size at a cellular growth cost loses its compensating benefit during unicellular growth, reducing the number of mutant founders entering the next multicellular phase. When those descendants form only one or a few groups, stochastic loss via genetic drift can eliminate the lineage before its net benefit is realized.

Prior work has largely conflated within-group relatedness with obligate vs. facultative multicellularity, and no experiment has decoupled these factors in a single model system. To bridge this gap, we engineered a *Saccharomyces cerevisiae* strain capable of switching between unicellular and clonal multicellular growth, allowing us to directly compare the evolutionary consequences of obligate versus facultative life cycles while holding genetic background constant. Over 192 days of experimental evolution, obligately multicellular populations consistently evolved larger clusters and rapidly acquired tetraploidy, a key adaptation for increased size that may carry a cellular growth cost. In contrast, populations with even brief unicellular phases were more constrained, with most remaining diploid and showing minimal increases in cluster size. Our theoretical analysis suggests that phase-dependent selection and stochastic loss interact to constrain rare multicellular adaptations. Unicellular growth interrupts selection for larger groups, while cellular growth costs of multicellular adaptations can reduce mutant abundance before the next bottleneck.

## Results

### Engineering a yeast strain capable of growing as single cells and clusters

Snowflake yeast are a model system of simple, undifferentiated multicellularity. They grow clonally, originating via mutations (e.g., *ace2/ace2*) that disrupt daughter-cell separation during mitosis^40^. The group reproduces via branch fracture, driven by the physical strain that cellular division places on cell-cell connections. While standard snowflake yeast are obligately multicellular, we engineered a strain capable of expressing varied life cycles, including obligately multicellular, unicellular, or facultatively multicellular cycles (Figure 1a). Specifically, we placed the transcription factor *ACE2* under the regulation of LexA-ER-AD^41^, a transcriptional activator controlled by exogenous β-estradiol (Figure 1b). When β-estradiol is present, *ACE2* is induced and cells grow as singles; when absent, *ACE2* remains inactive and cells form multicellular groups. By alternating between media with and without β-estradiol, the yeast experiences a facultative multicellular life cycle (Figure 1c,d; Supplementary Video 1).

**Figure 1.**
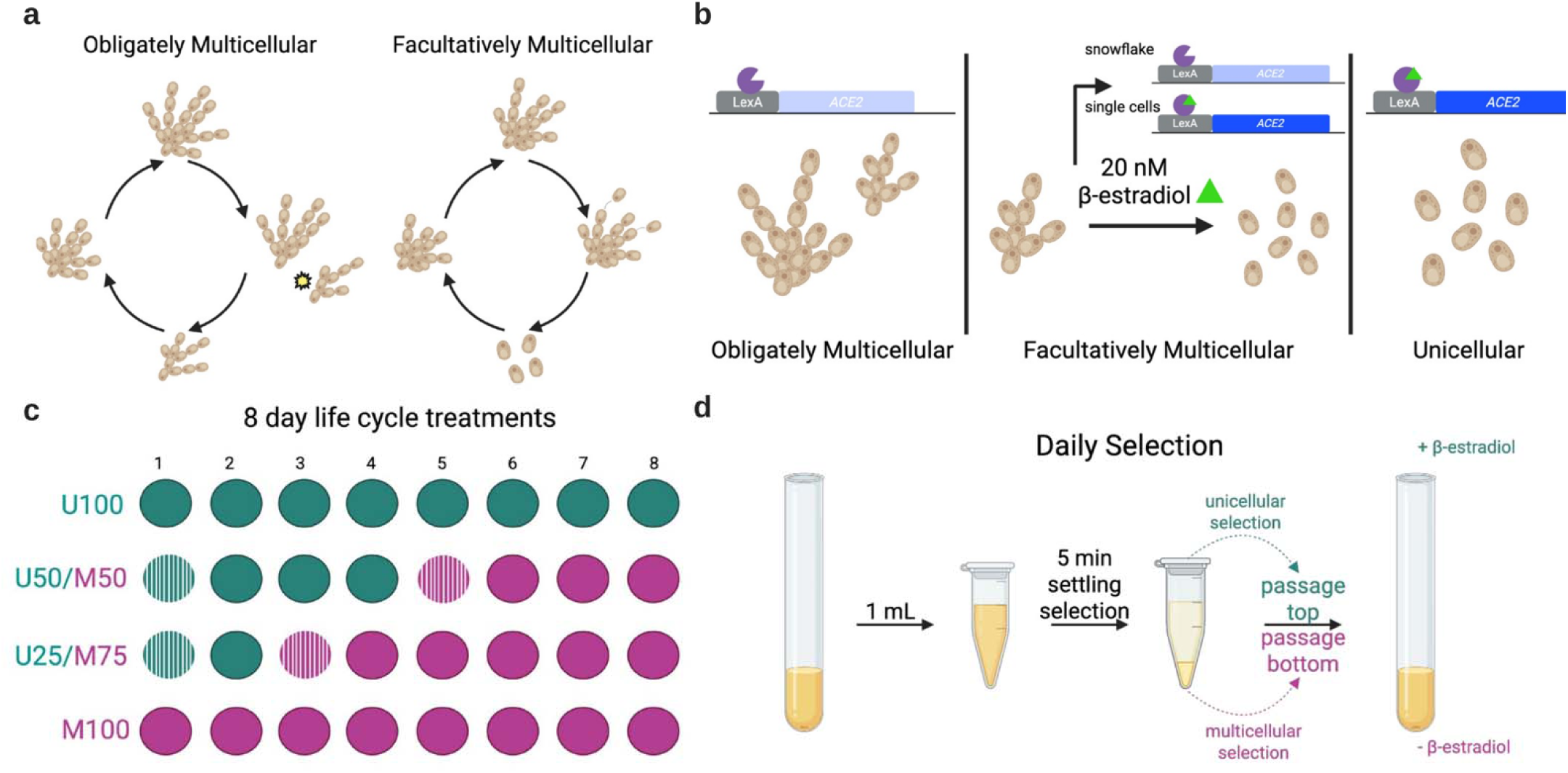
Engineering a facultatively multicellular strain of yeast. We created both obligate and facultative multicellular life cycles (a) by engineering a strain of *S. cerevisiae* in which *ACE2* is under exogenous regulation. We replaced the endogenous *ACE2* promoter with an artificial promoter driven by LexA*-ER-AD,* which is active only when the growth media contains β-estradiol. This strain grows as standard, obligately multicellular snowflake yeast when the media lacks β-estradiol (c, ‘M100’ life cycle), and is obligately unicellular when β-estradiol is always present (c, ‘U100’ life cycle). We also created two facultatively multicellular life cycles that vary in the proportion of time spent as clonally-developing snowflake yeast, or unicells (c, ‘U50/M50’ and ‘U25/M75’life cycles). We reinforced the change in growth form between unicellular and multicellular states with daily selection favoring that phenotype: during a unicellular phase, we selected cells from the supernatant after 5 minutes of settling selection (d), while during the multicellular phase, we selected from groups in the pellet.

### Experimental evolution of facultative vs. obligate multicellular life cycles

We examined four different life cycles: obligately unicellular (100% unicellular; ‘U100’), obligately multicellular (100% snowflake; ‘M100’), and two facultatively multicellular life cycles, one alternating equally between states (50% unicellular/50% multicellular; ‘U50/M50’), and one featuring a transient unicellular phase (25% unicellular/75% multicellular; ‘U25/M75’) (Figure 1c). We evolved five replicate populations of each life cycle (20 populations in total) over 24 eight-day periods. Each population underwent daily selection appropriate to its current life cycle phase. For the multicellular phase, we selected for larger, faster-settling clusters by allowing a brief settling period, then transferring clusters from the bottom of the culture tube (Figure 1d, Supplementary Figure 1A). This approach, the core selective regime of the Multicellularity Long Term Evolution Experiment (MuLTEE)^12,13,40,42^, has been shown to promote multicellular adaptation in snowflake yeast. During the unicellular phase, we selected cells from the top of the culture after settling, effectively removing any remaining clusters within a single round of selection (Figure 1d, Supplementary Figure 1B,1C). We relaxed selection during transition periods between each growth phase, by transferring 50 uL of culture to fresh medium without size-based selection for one day after switching media. To ensure comparable population size measurements, we transferred twice as much culture (100 µl) during the unicellular phase as during the multicellular phase, as the settled pellet contained higher cell density. We did not detect a difference in total generations among treatments across the eight- day cycle (Supplementary Figure 2). Populations were archived in glycerol stocks at -80 °C at the end of every eight-day cycle.

To confirm expected differences in population dynamics, we measured population size (as units of selection per culture) across all four life cycles (Figure 2). The obligately multicellular life cycle exhibited 11.2-fold lower population size than the unicellular life cycle (95% CI, 7.43–16.7), with geometric means across the eight-day cycle of 5.82 × 10^7^ clusters vs 6.49 × 10^8^ single cells at stationary phase (Holm-adjusted *p* = 0.0008). Both alternating life cycles displayed population sizes that corresponded to their current phenotypic state. The U50/M50 life cycle had a geometric mean of 7.38 × 10^8^ single cells during the first four days, declining to 4.94 × 10^7^ clusters during the last four days, a 14.9-fold difference (95% CI, 1.69– 132; Holm-adjusted *p* = 0.0333). Similarly, the U25/M75 life cycle went from 1.39 × 10^9^ single cells during the first two days to 5.45 × 10^7^ clusters during the remaining six days, a 25.5-fold difference (95% CI, 18.4–35.2; Holm-adjusted *p* = 0.0011). All three cultures in each facultative treatment had lower population sizes during multicellular growth, consistent with successful phenotypic switching. A sensitivity analysis allowing separate trends within each phase was less precise for U25/M75 (Supplementary Figure 3).

**Figure 2.**
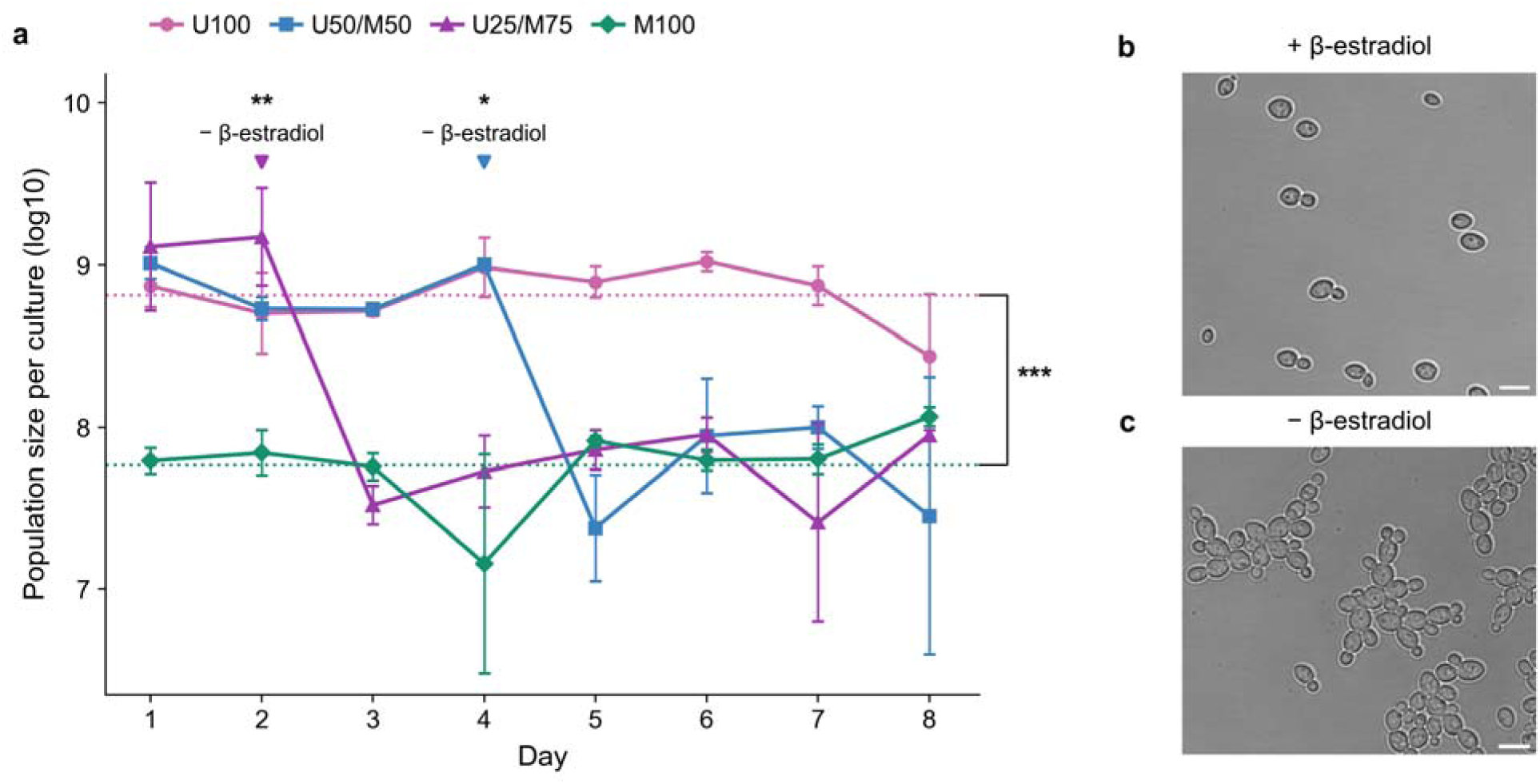
Population dynamics across the four life cycles. (a) Population size, measured as units of selection per culture, across the eight-day transfer regime. Points and error bars show mean ± SD of log_10_-transformed population sizes (*n* = 3 independent cultures per life cycle, each followed across all eight days). Dotted lines show the eight-day means of log_10_ population size for U100 and M100. Arrows mark the scheduled removal of β-estradiol. Asterisks indicate two- sided *p* values after Holm adjustment (\**p* < 0.05, \*\**p* < 0.01, \*\*\**p* < 0.001) for three culture- level comparisons: U100 versus M100 using Welch’s *t* test on eight-day culture means, and unicellular versus multicellular phase means within each facultative treatment using paired t tests. Phase comparisons use days 1–4 versus 5–8 for U50/M50 and days 1–2 versus 3–8 for U25/M75. Representative images show the facultative strain growing as single cells with β- estradiol (b) and as snowflake clusters without β-estradiol (c). Scale bars, 5 μm.

After 24 8-day cycles (192 days of evolution), cluster size differed among evolved multicellular treatments (one-way ANOVA, *F*_2,12_ = 4.08, *p* = 0.0445; Figure 3a). M100 populations had larger clusters than U50/M50 populations (Tukey’s HSD, *p* = 0.0373), while comparisons involving U25/M75 were not significant (both *p* ≥ 0.236). Mean cluster radius was 23.1% higher than the measured multicellular ancestor reference in M100, 10.4% higher in U25/M75 and 2.2% higher in U50/M50. The ancestor was excluded from statistical tests.

**Figure 3.**
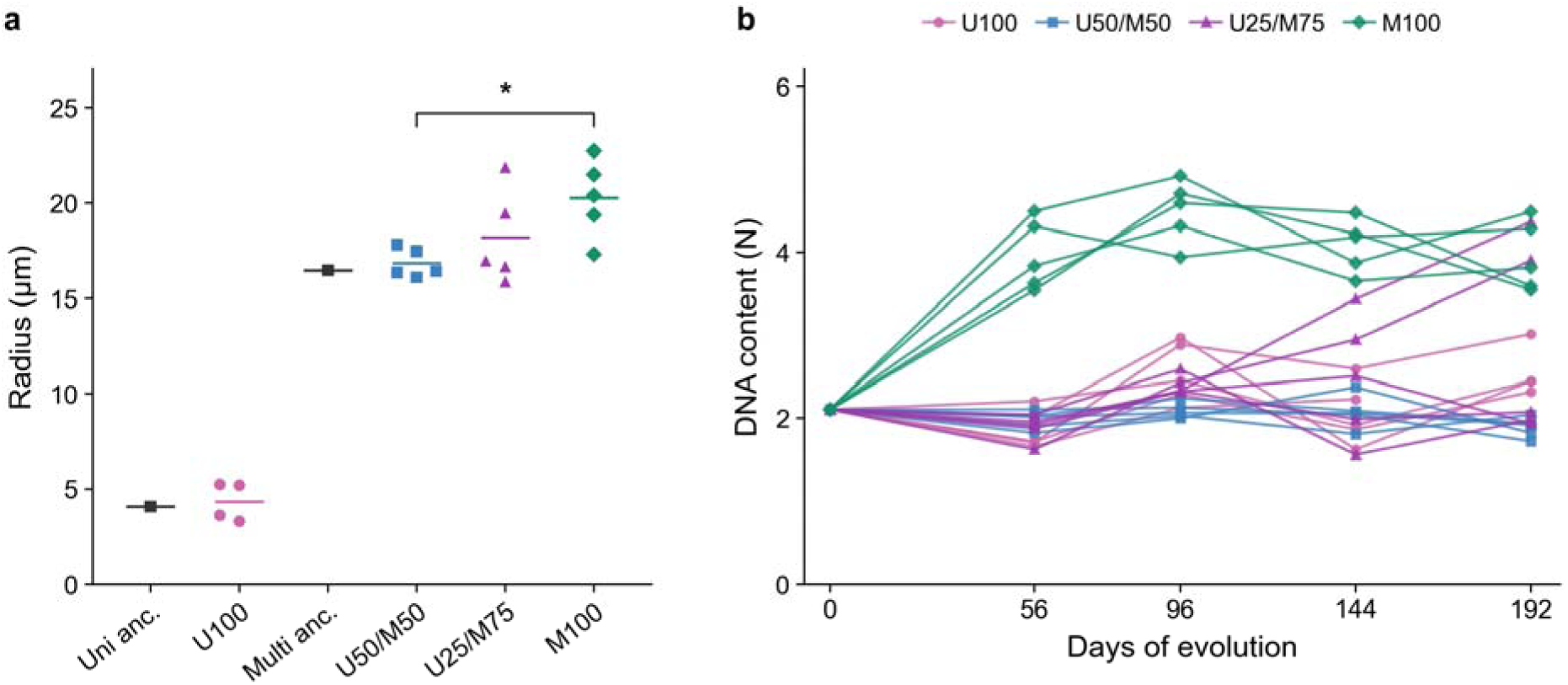
Obligate multicellularity promotes multicellular adaptation via whole genome duplication. (a) Cluster radius after 192 days of evolution. Colored points show population means (*n* = 5 per evolved multicellular treatment; *n* = 4 for U100); horizontal bars show treatment means. Black points show ancestor references. Statistical comparisons include only M100, U50/M50 and U25/M75; the bracket marks the M100 versus U50/M50 comparison (\**p* < 0.05 after Tukey adjustment). (b) DNA content analysis reveals that tetraploidy evolved in all five obligately multicellular populations, but was constrained in facultative life cycles, with only two of five U25/M75 populations evolving tetraploidy, while all U50/M50 populations remained diploid throughout. Each point in b represents a population mean.

The earliest, most convergent adaptation underlying larger group size in the MuLTEE is whole genome duplication, resulting in tetraploidy^13^, which arose independently in all 10 anaerobic and mixotrophic populations within the first 200 days. Tetraploidy increases cluster size (and thus multicellular fitness) by increasing both cell size and cellular aspect ratio^13^. Here, we find that tetraploidy emerged in all five obligately multicellular populations by day 56, and persisted for the full duration of the experiment (192 days). In contrast, facultative populations showed markedly different outcomes: only two of five U25/M75 populations evolved tetraploidy by day 192, while all U50/M50 populations remained diploid throughout (Figure 3b). These results suggest that the presence of a unicellular growth phase, even a relatively brief one, may constrain the evolution of novel multicellular traits.

### Establishment barriers limit the evolution of tetraploidy in facultative populations

The observed differences in cluster size and ploidy across our four life cycles raise a fundamental question about the underlying evolutionary mechanisms. The absence of tetraploidy in most facultative populations could reflect two scenarios: tetraploidy may carry a net fitness disadvantage due to costs imposed during unicellular phases, or facultative life cycles may be unable to fix a net beneficial mutation because of establishment barriers created by the single cell phase. To ask whether tetraploidy is expected to increase across each facultative life cycle, we used phase-specific fitness estimates to simulate its frequency from a 50% starting point.

We estimated unicellular fitness by competing 2N and 4N strains separately against the same GFP competitor (*r* = −0.074 per day, SD = 0.364, *n* = 3 replicate contrasts). The mean was similar to that for multicellular growth without settling selection (*r* = −0.050 per day, SD = 0.161, *n* = 4)^13^. Published mixotrophic multicellular competitions with settling selection showed a fitness advantage for tetraploids (*r* = 0.471 per day, SD = 0.105, *n* = 4)^13^. Combining our estimate of the fitness cost of tetraploidy in a unicellular state with its multicellular advantage during settling selection in a Wright-Fisher simulation predicted a net benefit of tetraploidy across both facultative life cycles (Figure 4a). Starting at equal frequency, tetraploids declined during unicellular phases but gained more during multicellular phases, approaching fixation over two successive 8 day cycles. So why was it so much more difficult for tetraploidy to evolve in the facultative life cycles than in the obligately multicellular treatment?

**Figure 4.**
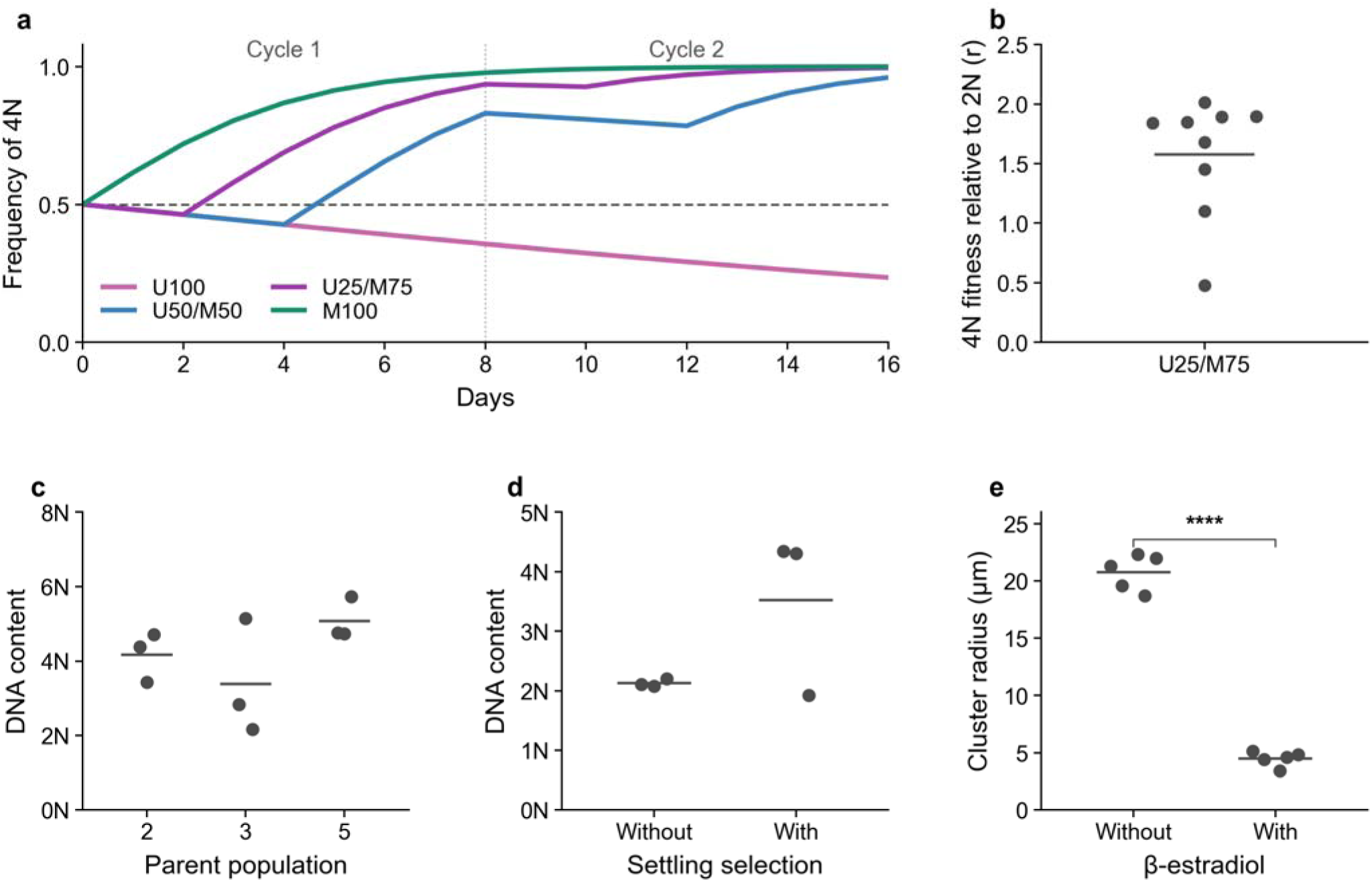
Tetraploidy is strongly beneficial in facultative life cycles, even though it rarely evolved within them. (a) Wright-Fisher simulations using our unicellular selection rate and the published multicellular settling rate^13^. Tetraploids start at 50% frequency. Lines show means of 1,000 simulations per regime across two eight-day cycles, separated by the vertical dotted line. Tetraploids increased across successive cycles in all three regimes containing a multicellular phase. We tested this empirically (b), competing tetraploids and diploids at equal initial ratios across one U25/M75 life cycle in nine independent competition cultures, spanning all pairings of three diploid and three tetraploid isolates. As with the simulation, tetraploidy was strongly adaptive in this life cycle. We performed a series of controls to rule out alternative explanations. (c) U25/M75 populations that remained diploid for the full 192-day experiment evolved tetraploidy in 7/9 replicate populations after just 48 days of obligately multicellular evolution, demonstrating that these lineages retained the capacity for whole genome duplication (WGD). (d) Tetraploidy did not evolve in the absence of settling selection, confirming that WGD is specific adaptation to size-based selection. (e) Tetraploids retained the capacity for unicellular growth with β-estradiol (two-sided paired *t*-test, *n* = 5 populations measured with and without β- estradiol; \*\*\*\**p* < 0.0001). Points show population means; lines show condition means.

Facultative life cycles may impose an establishment barrier on tetraploidy. During unicellular growth, selection for larger groups is absent, and any cellular growth cost can reduce the number of tetraploid descendants entering subsequent transfers. Multicellular growth can then leave those descendants in only one or a few groups, increasing their risk of loss at a daily bottleneck. To test whether tetraploidy remained beneficial when introduced at equal frequency, we competed isogenic tetraploid and diploid strains across one U25/M75 life cycle. Consistent with our model, tetraploidy was strongly beneficial across the full 8-day life cycle when competing at equal frequency (*r* = 1.58; one sample *t*-test, *t* = 9.483, *n* = 9 competition cultures, *p* < 0.0001; Figure 4b). This establishes a net benefit at the tested starting frequency, consistent with an establishment barrier acting when tetraploidy is rare.

To confirm that the absence of tetraploidy in most facultative populations reflected life cycle dynamics rather than an inherent inability to undergo whole genome duplication, we conducted a follow-up experiment with the U25/M75 populations. We selected the three populations that remained diploid throughout the evolution experiment and re-evolved nine replicate populations (three of each) under obligately multicellular conditions. Within 48 days, seven out of the nine populations evolved tetraploidy (Figure 4c). This confirms that failure to evolve tetraploidy was not due to an inherent limitation on genome duplication, but to the constraining effect of the unicellular growth phase.

We conducted two further control experiments to rule out alternative explanations for our results. We tested (1) whether tetraploidy evolves as a consequence of obligate multicellular growth rather than as a specific adaptation to size-based selection, and (2) whether evolved tetraploid populations are physiologically constrained from transitioning between growth modes, a requirement for success in facultative populations. First, we re-evolved obligately multicellular strains with and without selection for larger group size. After 40 passages, tetraploidy evolved only in populations under size selection, confirming that genome duplication is a specific adaptation to size-based selection rather than an inevitable consequence of multicellular growth (Figure 4d). Second, we tested whether tetraploid populations retained the capacity to switch between growth modes by growing them with β-estradiol. If tetraploid cells could not transition to unicellular growth, they would be unable to complete the facultative life cycle, providing an alternative explanation for why tetraploidy rarely evolved in these populations. All five populations successfully transitioned to unicellular growth, confirming that tetraploid cells remain capable of alternating between growth modes (two-sided paired *t*-test, *t* = 19.77, *n* = 5 populations, *p* < 0.0001; Figure 4e). Taken together, these controls suggest that the absence of tetraploidy in most facultative populations likely reflects life cycle-dependent evolutionary dynamics rather than physiological or experimental constraints.

### Theoretical framework for life-cycle driven establishment barriers

To understand how life cycle structure affects the establishment of new mutations, we first considered a neutral bottleneck model, building on the framework of Pentz et al.^29^. Populations grow *f*-fold before a random sample returns them to *N* cells. We compared sampling individual cells with sampling equally sized groups (Supplementary Note 1), isolating the effect of organizing mutant descendants into different numbers of units of selection.

A mutant confined to one unfragmented group has only one chance to survive the bottleneck. If one in every 32 groups is retained at random, that group has a 1/32 chance of survival and a 31/32 (96.9%) chance of being lost. This is true whether the group contains 10 or 100 cells: cell division increases the number of mutant cells without creating additional units of selection. In unicellular growth, each cell division creates another separately sampled cell, reducing the probability that all mutant descendants are lost.

Alternating life cycles repeatedly change this relationship between cell number and the number of units of selection. Dispersal releases the cells in a surviving group into separately sampled units, and subsequent unicellular growth can further reduce loss through drift. In the reverse transition, each surviving cell founds a clonal group. If a rare mutant is represented by only one founder, its descendants remain a single unit of selection until that group fragments. Growth within the group therefore provides no additional protection against a neutral bottleneck. Cellular growth costs can also reduce the number of mutant founders reaching the multicellular phase, limiting opportunities for later group-level benefits to be realized.

To examine how fitness effects interact with these bottlenecks, we simulated mutation survival across our four life cycles for eight rounds of growth and selection, beginning with a single mutant cell (Methods). Populations began each round with 10^6^ cells, expanded 32-fold, and were then reduced by sampling cells or groups, depending on the life cycle. We varied the mutant’s exponential growth-rate multiplier and group-selection weight while holding total population expansion constant. For each parameter combination and life cycle, we averaged survival across three batches of 1,000 simulations. In the purely unicellular life cycle (U100), survival depended on cellular growth rate alone, because group-level selection was absent (Figure 5b). In the obligately multicellular life cycle (M100), both faster growth and increased group-selection weight improved survival (Figure 5a). Mutations with negative pleiotropy, conferring multicellular benefits at a cellular growth cost, tended to survive when their group- level benefits outweighed cellular growth costs. These simulations measure eight-round survival, a prerequisite for surviving the establishment barrier, rather than eventual fixation.

**Figure 5.**
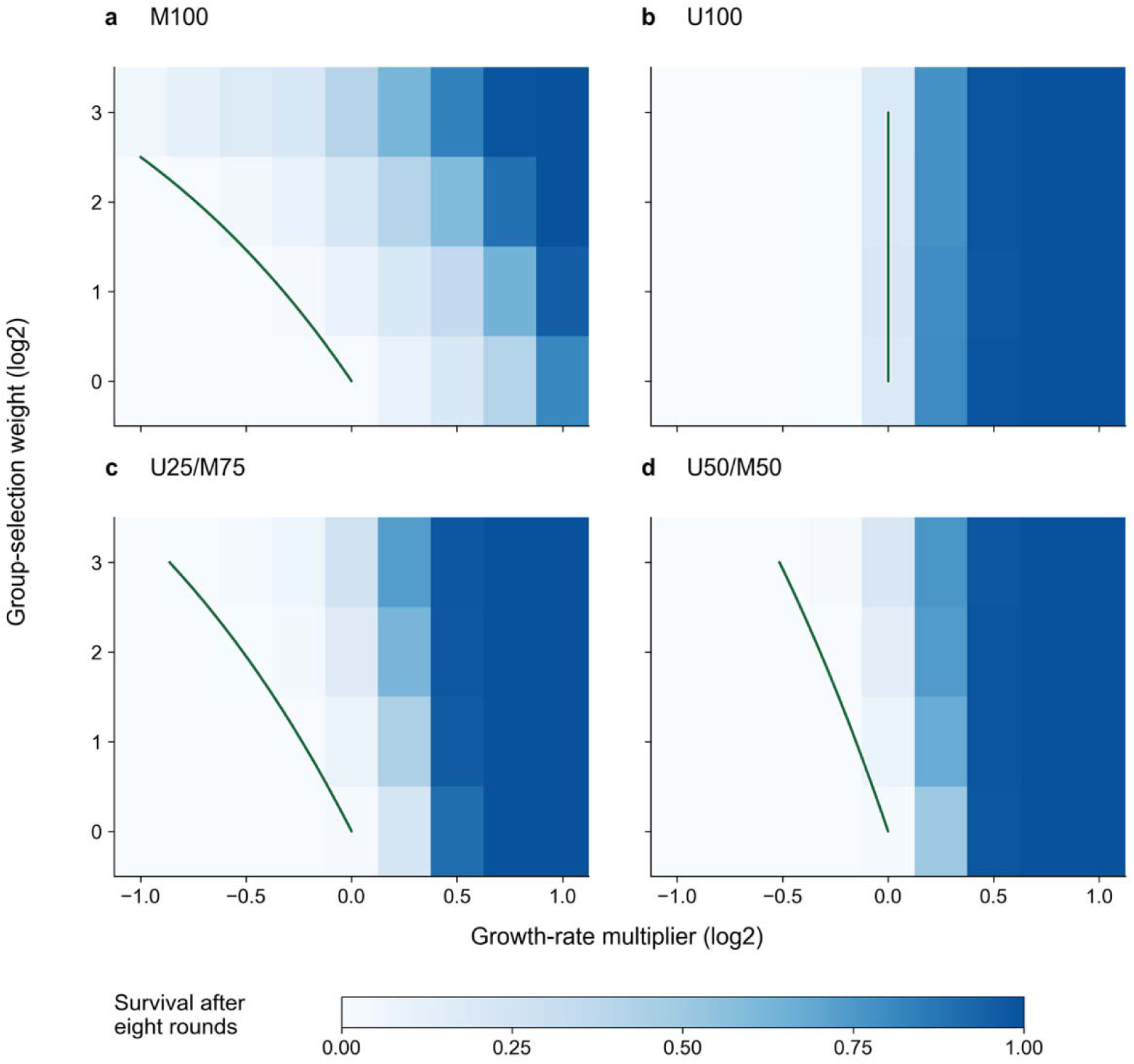
Life cycle structure shapes the survival of new mutations. Panels show M100 (a), U100 (b), U25/M75 (c) and U50/M50 (d). Shading gives the mean fraction of mutant lineage surviving eight rounds of growth and selection across three batches of 1,000 simulations per parameter combination. Each lineage began as a single mutant cell. The axes show log_2_ values of the mutant’s exponential growth-rate multiplier *g* and group-selection weight *w*, relative to the ancestor. Green lines mark relative cycle fitness *W* = 1 in the deterministic rare-mutant limit; mutations above the curves (a,c,d), or to the right of the vertical line (b), have *W* > 1. The multicellular isoclines are curved on the plotted growth-rate axis (Supplementary Note 1, equations 6-8). Mutants with a net fitness benefit can nevertheless have low survival when cellular growth costs continue through rounds without selection for larger groups.

Both facultative life cycles displayed intermediate survival patterns, with a pronounced dependence on cellular growth rate and a weaker response to group-selection weight (Figure 5c,d). Growth effects act in every round, whereas group-level benefits act only during multicellular rounds. The green isoclines mark fitness equal to the ancestor in the deterministic rare-mutant limit (Supplementary Note 1, equations 6-8). Mutations above these curves are net beneficial, yet those with cellular growth costs often had low survival in facultative life cycles. Early bottlenecks can therefore prevent rare lineages from realizing their later group-level benefits. Hierarchical clustering of the 36 mean survival counts per life cycle placed both facultative cycles closer to U100 than to M100 (Supplementary Figure 4). Thus, even brief unicellular phases can constrain the establishment of mutations that improve multicellular performance.

## Discussion

Life cycle structure shapes the selective landscape of nascent multicellular organisms, governing the balance of selection at the cellular and collective level^16,29,31,43^. While extensive prior work has focused on the role of within-group genetic relatedness in shaping multicellular evolution^5,22,24,25^, little work has examined how obligate versus facultative multicellular life cycles shape evolutionary outcomes independent of relatedness. Our results demonstrate that even brief periods of unicellularity can fundamentally constrain multicellular adaptation, preventing the establishment of beneficial mutations that would otherwise be highly advantageous across the complete life cycle.

In our system, strongly beneficial mutations like tetraploidy were considerably less likely to evolve in facultative life cycles, despite providing substantial net fitness benefits. We can infer that mutations creating tetraploidy occur readily in all treatments, as we observed convergent evolution five times in the obligately multicellular populations in this experiment (Figure 3b), in 10 obligately multicellular populations of the MuLTEE^13^, and in 7/9 replicate populations when previously-diploid U25/M75 populations were re-evolved under obligately multicellular conditions for just 48 days (Figure 4c). Why aren’t these overwhelmingly beneficial mutations (Figure 4a) being driven to fixation by selection in facultative multicellular life cycles?

Our results show how life cycles can impose establishment barriers to multicellular adaptations. Group formation packages many cells into fewer units of selection^36,38^. When a mutant is confined to one unfragmented group, cell division increases its abundance without increasing the number of units that can survive a bottleneck. Dispersal restores cell-level sampling, but growth costs during the unicellular phase can reduce the mutant founders available to establish new groups. In our simulations, mutations with multicellular benefits and cellular growth costs had lower survival in facultative than in obligately multicellular life cycles (Figure 5a,c,d). Even mutations with a positive fitness effect across a complete facultative cycle were frequently lost within eight rounds, showing how early loss can prevent a lineage from realizing

its later benefits. Such antagonistic pleiotropy is likely common during the evolution of multicellularity, as cellular traits underlying multicellular innovations often impose fitness costs in unicellular contexts^21,23,44–47^.

These findings provide insight into one of the most striking results from comparative biology: ’complex’ multicellularity has only evolved five times out of the 45 independent origins of multicellularity^1^, in all cases within lineages which are both obligately multicellular and develop clonally^19^. Both traits may play a central role in the evolution of increasingly complex multicellular organization, with clonality helping solve the problems of within-group genetic conflict^9,48,49^ and multicellular heritability^27^, while obligate multicellularity circumvents establishment barriers for novel multicellular traits. Clonality and obligate multicellularity covary in commonly studied life cycles (’staying together’ after cell division yields clonal, obligately multicellular groups, while ’coming together’ yields chimeric, facultative groups), but this is not a hard requirement. Notably, many organisms that facultatively form clonal multicellular groups (and should thus have little scope for within-group conflict), including choanoflagellates^33^, *Chlamydomonas priscuii* and *Chlamydomonas raudensis*^32^, and cyanobacteria^34^, have remained comparatively simple despite hundreds of millions of years of evolution^11,50,51^. While multiple factors likely contribute to this pattern, the establishment barriers imposed by facultative life cycles may play a role.

A large body of theoretical and empirical work has established that clonal development is central to the evolution of complex multicellularity, maintaining high within-group relatedness and thereby suppressing social conflict and aligning the fitness interests of cells within groups^5,22,48,52,53^. Our results complement these findings by revealing that obligate multicellular life cycles provide an additional, independent benefit: even within clonal groups where social conflict is not a factor, facultative life cycles still impose establishment barriers that stem from the population genetic consequences of group formation. Together, these results suggest that clonality and obligate multicellularity facilitate the evolution of increasingly complex multicellularity through at least two distinct mechanisms, the former suppressing within-group conflict and the latter removing establishment barriers for novel multicellular adaptations. Because any evolutionary transition in individuality necessarily collapses many lower-level units of selection into fewer collective-level units, the constraints we identify may extend beyond multicellularity to other ETIs.

## Methods

### Strains and Media

We engineered *S. cerevisiae* to switch between unicellular and snowflake multicellular growth using the LexA-ER-AD inducible system^41^. LexA-ER-AD was integrated at the *HO* locus, and the native *ACE2* promoter was replaced with a synthetic promoter containing a single LexA binding site. In this strain, addition of β-estradiol induces *ACE2* transcription, driving expression of downstream cell separation enzymes and restoring unicellular growth.

All experiments were performed in Yeast Peptone Dextrose [10g yeast extract, 20g peptone, and 20g dextrose per liter] medium. All replicate populations were grown at 30 °C with 225 rpm shaking. To induce unicellular growth, YPD was supplemented with β-estradiol to a final concentration of 20 nM^54^. β-estradiol stock solution was prepared by dissolving 0.04 g β- estradiol in 1 mL ethanol, then diluting in H_2_O to 10^-5^ M concentration. For experimental use, 20 uL of stock was added per 10 mL fresh YPD medium.

### Inducible strains

yAP201: Y55HD background, ace2Δ:KanMX4/ace2Δ:KanMX4, ho:PACT1(-1-520)-LexA-ER- haB42-TCYC1 prACE2Δ::PTEF-HygMX-TTEF-insul-(lexA-box)1-PminCYC1

### Plasmids

pAP010: Expression of *LexA-ER-B42* β-estradiol inducible system under *ACT1* promoter

pAP012: Expression of inducible promoter containing 1 lexA box

### Oligonucleotides used for replacing the *ACE2* promoter with inducible promoter

ACE2_est_F: AAGAAATAACTAAAGAAATCTATAGGACCAAAAACGGTGTTAATACAATCCGAGAGCTTGCCTTGTCC

ACE2_est_R: CTTTCGCGAAGCCTGAGGGATTTATATACCACGGATCTACAACGTTATCCATAAGCTTGATATCGAATTCCTG

Strains and plasmids were adapted from Barrere et al.^54^ and Ottoz et al.^41^.

### Evolution experiment

We evolved five replicate populations under each of four life cycle treatments: purely unicellular (U100), obligately multicellular (M100), and two facultatively multicellular treatments that alternated between growth modes, one with equal time in each phase (U50/M50) and one with a shorter unicellular phase (U25/M75). To establish ancestral populations, we grew a single isogenic clone overnight, then transferred 100 µL aliquots into fresh medium lacking β-estradiol (multicellular growth) or supplemented with β-estradiol (unicellular growth). The U100 treatment was initiated from the unicellular condition, while the U50/M50, U25/M75, and M100 treatments were initiated from the multicellular condition.

Every 24 h, each population was subjected to stage-specific daily selection. During multicellular growth, we used settling selection to select for larger cluster size. After 24 h of growth, we transferred 1 mL of culture into 1.5 mL Eppendorf tubes and let them settle on the bench for 5 min. The top 950 µL of culture was discarded, and the bottom 50 µL was transferred into 10 mL of fresh medium for the next round of growth and selection. During unicellular growth we followed the same protocol as above, but transferred 100 µL from the top of the settled culture rather than 50 µL from the pellet, as the lower cell density in the supernatant requires a larger volume to transfer comparable biomass. For the facultatively multicellular life cycles, we relaxed selection during transitionary periods between each growth phase (during which unicells grow into new clusters and clusters shed unicells, establishing the new population state) by transferring 50 µL of culture without size-based selection. Each population evolved continuously for 192 days, with frozen glycerol stocks archived at -80 °C every 8 days (24 total life cycles per population). At day 192, we excluded U100 Rep 3 because its cells clumped and grew as groups rather than individual cells; its earlier ploidy measurements are retained in Figure 3b.

### Evaluating population dynamics

We measured the population size of all four treatments across an 8-day life cycle, with three replicates per treatment. Each population was initially grown for 24 h, followed by one additional transfer to stabilize populations prior to measurement. We estimated population size (number of clusters or cells, depending on whether the population was multicellular or unicellular, respectively) for each timepoint in 24-well plates by preparing a 10^3^-fold dilution to minimize cluster overlap, and imaged the whole well with a Nikon Eclipse Ti microscope with 40x magnification. This procedure was repeated at 24 h intervals throughout the 8-day experiment, with samples collected at both initial and final time points.

To account for repeated measurements, we treated each culture as the independent replicate and analysed log_10_-transformed population sizes. We compared the eight-day mean for each U100 and M100 culture using a two-sided Welch’s *t* test (three cultures per treatment; *t* = 18.48, *df* = 3.13). For each facultative culture, we fitted a piecewise-constant regression with separate unicellular and multicellular phase means, then tested the three pre-minus-post coefficients against zero using a two-sided one-sample *t* test. This is equivalent to a paired *t* test of the culture-specific phase means (U50/M50: *t* = 5.34, *df* = 2; U25/M75: *t* = 42.97, *df* = 2). We applied Holm adjustment across these three comparisons. Geometric-mean ratios and pointwise 95% confidence intervals were obtained by back-transforming the log_10_ contrasts and their *t*- based intervals; confidence intervals were not adjusted for multiple comparisons.

As a sensitivity analysis, we fitted log_10_(*N*) = *a* + *b*(day − *k*) + *cI* + *d*(day − *k*)*I* separately to each facultative culture, where *k* is the first multicellular measurement (day 5 for U50/M50 and day 3 for U25/M75) and *I* is 1 from day *k* onward and 0 otherwise. We tested the mean of the three −*c* coefficients against zero with a two-sided *t* test (*df* = 2), reporting unadjusted *p* values and 95% confidence intervals (Supplementary Figure 3). This contrast measures separation of the extrapolated pre-switch and post-switch trends at day *k*. U25/M75 has only two pre-switch measurements per culture, limiting precision. These analyses assume approximately normal culture-level summaries or contrasts, which cannot be assessed reliably with three cultures, and the fixed switching schedule links phase to elapsed time. All 96 observations were retained. Supplementary Code 3 reproduces the analyses and figures.

### Measuring cluster size

Cluster size distributions of ancestor and evolved populations were measured via microscopy and analyzed using ImageJ. We revived each population and inoculated strains by transferring 100 uL into 10 mL of YPD media and grew them for 24 hours at 30 . We then transferred 100 uL into fresh 10 mL of YPD and grew them for an additional 24 hours to stabilize the population. Cultures were diluted 1,000-fold prior to imaging to minimize cluster overlap. 300 uL of diluted sample was transferred to individual wells of 24-well plates. Images were captured using a Nikon Eclipse Ti inverted microscope with 40x magnification. For each well, we scanned the entire well by capturing a 5 x 8 grid of images, with 10% overlapping stitching. We used a custom ImageJ analysis pipeline to measure the cross-sectional area of individual clusters.

To compare cluster size among evolved multicellular treatments, we used one mean radius per independently evolved population (five populations each for U50/M50, U25/M75 and M100). We performed a one-way ANOVA followed by Tukey’s HSD across all three pairwise comparisons at a family-wise α = 0.05, with simultaneous 95% confidence intervals for mean differences. Ancestor values and U100 populations were excluded from these tests. As a sensitivity analysis allowing unequal variances, we used Welch’s ANOVA (*F*_2,6.35_ = 5.77, *p* = 0.0373) and Games–Howell comparisons. These gave the same qualitative result: M100 differed from U50/M50 (adjusted *p* = 0.0388), while the other pairs did not resolve differences. Supplementary Code 4 reproduces the analysis and Figure 3a.

To test whether evolved tetraploid populations retained the capacity for unicellular growth, we measured mean cluster radius in each of five independently evolved populations with and without β-estradiol (Figure 4e). We compared conditions using a two-sided paired *t*-test on the five within-population differences in mean radius.

### Ploidy measurement

Ploidy levels were estimated using an imaging-based approach developed by Tong et al.^13^ for multicellular *S. cerevisiae,* with minor modifications as described below. Briefly, we used microscopy to image propidium-iodide-stained clusters to quantify the fluorescent intensity of nuclei.

For each experiment, populations were grown overnight in YPD media (supplemented with -estradiol for U100 populations) along with engineered 2N and 4N control strains. Cultures were then transferred to fresh YPD media and grown to mid-log phase. For each population, 250 ul of culture was transferred into microcentrifuge tubes, pelleted by centrifugation (5000 g, 1 min), washed with 1 mL H_2_O, and fixed using 70% ethanol for 2 hours at room temperature with rotation. Fixed cells were washed twice with 50 mM sodium citrate, treated with RNase A (0.5 mg/mL) at 37 °C for 2 hours, and stained overnight with propidium iodide (5 µl of 1 mg/mL stock) at 30 °C in the dark with rotation.

For imaging, 5 μl of stained clusters were flattened onto a microscope slide. Images were captured at 200x magnification using a Nikon Eclipse Ti inverted microscope with identical settings to Tong et al.^13^: red channel (600 ms exposure, 2.2x gain) and brightfield channel (100ms exposure, 4.1x gain). Approximately 8-12 FOV were imaged per sample. Image analysis was performed in ImageJ (v1.54f) to quantify propidium iodide fluorescence intensity.

For U100 populations, which consisted of single cells rather than multicellular clusters, we used a lower size cutoff for minimum cluster area during nuclear segmentation to account for the smaller cell size, but otherwise applied the same analysis pipeline. DNA content was estimated by normalizing to the haploid genome calculated from the 2N and 4N control strains. Statistical analysis was performed in R(v4.4.1), with DNA content estimated based on the G1- phase peak intensities.

### Ploidy competition experiment

We competed isogenic 2N and 4N strains^13^ across an eight-day U25/M75 life cycle. We ran nine independent competition cultures, pairing each of three diploid isolates with each of three tetraploid isolates. We used a one-sample *t*-test across cultures, restricting inference to the tested isolates. Individual clusters were classified as diploid (2N) or tetraploid (4N) based on their estimated DNA content using the imaging-based ploidy measurement described above.

### Re-evolution under obligate multicellularity

We founded three descendant cultures from each of U25/M75 populations 2, 3 and 5 (nine cultures in total), and evolved them under obligately multicellular conditions for 48 days. We estimated ploidy using the imaging-based assay described above (Figure 4c).

### Mathematical modeling of evolutionary dynamics

We simulated the survival of a single mutant lineage through eight rounds of growth and selection. Populations began each round at 10^6^ cells and expanded 32-fold before sampling cells or groups. Growth followed for residents and for mutants, where *g* is the exponential growth-rate multiplier. We varied log_2_ *g* from -1 to 1 in increments of 0.25, keeping total expansion fixed. Mutant groups received selection weights *w* of 1, 2, 4 or 8. In multicellular rounds, we divided mutant cell number by group size (20 cells), rounding upward to obtain mutant group number. Cell sampling and unweighted group sampling used hypergeometric draws or a normal approximation. Weighted group sampling used Poisson or binomial approximations based on the weighted mutant-group frequency, with a deterministic branch when the expected mutant fraction among selected groups exceeded 0.95. The code specifies the sampling thresholds. Transitions to unicellular growth dispersed groups into individual cells; transitions to multicellular growth restored group-based sampling after growth. We used eight multicellular rounds for M100, eight unicellular rounds for U100, two unicellular followed by six multicellular rounds for U25/M75, and four of each for U50/M50. For each parameter combination and life cycle, we ran three batches of 1,000 simulations and plotted the mean fraction retaining at least one mutant after round eight. This endpoint does not establish fixation or guaranteed future survival. We calculated the fitness isoclines separately in the deterministic rare-mutant limit, combining the growth effect across all eight rounds with the group-selection effect in multicellular rounds (Supplementary Note 1, equations 6-8). This calculation omits stochastic loss, discrete group-count rounding and finite-frequency effects.

### Estimation of phase-specific fitness

For the unicellular estimate, we measured 2N and 4N strains separately against the same GFP competitor. For each ploidy, we averaged four GFP-fraction measurements at day 0 and four measurements from each of three replicate competitions at day 1. The day-0 mean was shared across the three replicates within each ploidy. For each competition, we calculated the change in the natural log of the non-GFP:GFP ratio. We then subtracted the diploid rate from the tetraploid rate for corresponding replicate indices, giving three estimates of the tetraploid fitness effect per day. We report their mean and sample standard deviation.

For the multicellular estimate, we used the four mixotrophic competitions with settling selection reported in Figure 3g of Tong et al.^13^ These competitions directly compared engineered diploid and tetraploid snowflake yeast over three days, estimating tetraploid frequency from the fraction of total cluster volume. We calculated each daily selection rate as the day-0 to day-3 change in the natural log of the tetraploid:diploid ratio, divided by three. This estimate includes growth and settling selection; no additional growth correction was added. We used the four published mixotrophic competitions without settling selection only for the descriptive comparison in the Results. The assays differed in frequency measurement and transfer protocol; only the settling-treatment mean entered the model as the multicellular rate. The selected inputs and their calculations are provided in the figure4_fitness_inputs sheet of Supplementary Data.

### Wright Fisher model simulation

We developed a stochastic Wright-Fisher model to simulate competition between diploid and tetraploid strains under each life cycle regime. Populations were initialized at a 1:1 ratio with a carrying capacity of 10 cells. Each day, we added the phase-specific selection rate to the log odds of tetraploidy and then sampled 1/32 of the population by binomial sampling. Unicellular phases sampled individual cells, while multicellular phases sampled clonal groups of ten cells. We used the unrounded means from Supplementary Data (approximately −0.073773 and 0.471286 per day for unicellular and multicellular phases, respectively). These rates describe the net fitness effects of each assay, including settling selection for the multicellular estimate. We ran 1,000 simulations per regime over two successive eight-day life cycles (Figure 4a). Rates were constant within each phase; the model did not separately represent the experimentally relaxed selection on transition days. Loss and fixation were absorbing states. Source data and simulation code are provided in the supplementary files.

### Time lapse microscopy

We used time-lapse microscopy to examine the transition from multicellular growth to unicellular growth in the engineered alternating strain. A single isolate was grown for 24 h in 10 mL YPD at 30 °C with 225 rpm shaking. On the next day, 1 mL of sample was transferred to 10 mL of fresh media for 4 h of growth so that the strain can reach an exponential growth phase.

1mL of sample was centrifuged, supernatant was removed and resuspended in YPD media. The sample settled on the bench for 5 min, and the bottom 100 uL of culture was transferred into a new 1.5 mL Eppendorf tube, with 900 uL of YPD supplemented with β-estradiol. 0.5 uL of sample droplets was placed into an 8 well chamber slide. To limit evaporation, 5 uL of H_2_O was placed on the walls of the chamber and the chamber was sealed with parafilm. Images were acquired at 15-minute intervals (Supplementary Video 1).

## AI assistance

We used Codex (OpenAI) to assist with code translation, figure preparation and final code review (numerical outputs from simulations were checked against source data, analytical expectations and archived simulation results).

## Data availability

Population-size measurements, cluster-radius summaries, ploidy trajectories, competition estimates and archived simulation outputs are provided in Supplementary Data (Supplementary_Data.xlsx). The Supplementary Information Guide describes the worksheets and identifies the figures they support.

## Code availability

Python code for the Wright–Fisher model and the mutant-survival model is provided in Supplementary Codes 1 and 2, respectively. Supplementary Code 2 also includes the original

MATLAB implementation and scripts for plotting survival probabilities and fitness isoclines. Supplementary Codes 3, 4 and 5 provide the population-size analyses, cluster-radius analyses and mutant-survival clustering, respectively. Each package includes instructions, software dependencies and numerical reference outputs.

## Supporting information

Multi to Uni Timelapse

Supplementary Wright Fisher Model

Supplementary Mathematical Model

Supplementary Population Size

Supplementary Cluster Radius

Supplementary Survival Clustering

## Acknowledgements

Figure 1 was created using BioRender.com.

## Author contributions

A.P. and W.C.R. conceived and designed the study. A.P. performed the experiments with assistance from A.B. and J.C. A.P. developed the Wright–Fisher simulation. E.L. developed the analytical model and performed the mutant-survival simulations. A.P. and W.C.R. analyzed the data and wrote the manuscript. W.C.R. supervised the project. A.P. and W.C.R. obtained funding.

## Additional information

Supplementary information accompanies this paper.

Correspondence and requests for materials should be addressed to William C. Ratcliff.

