## Supplementary Figure for "Obligate multicellularity circumvents population genetic barriers to collective-level adaptation"

**Supplementary Figures**

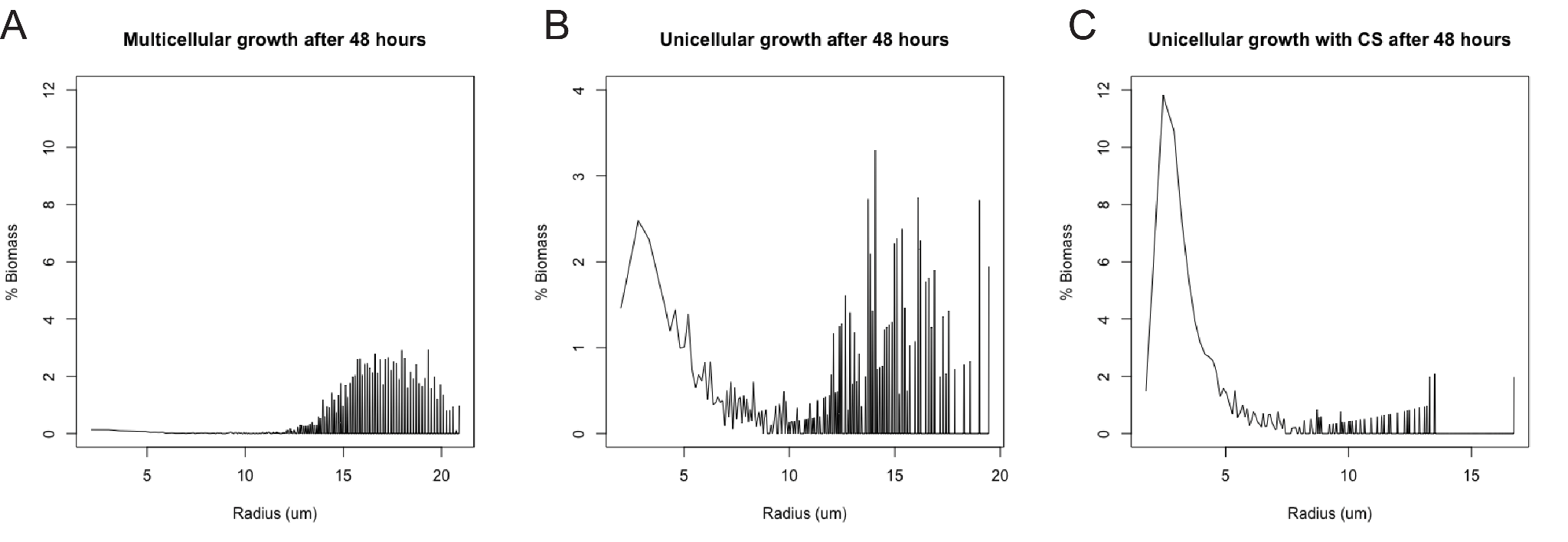

**Supplementary Figure 1. Size-based counter-selection is necessary to achieve predominantly unicellular populations after β-estradiol induction.** After 48 h of multicellular growth, cluster size peaked at ~17 µm (A). Induction with β-estradiol alone was insufficient to fully disperse clusters, leaving a substantial fraction of groups intact (B). A counter-selection (CS) protocol against groups yielded a predominantly unicellular population, with individual cell size peaking at ~3 µm (C).

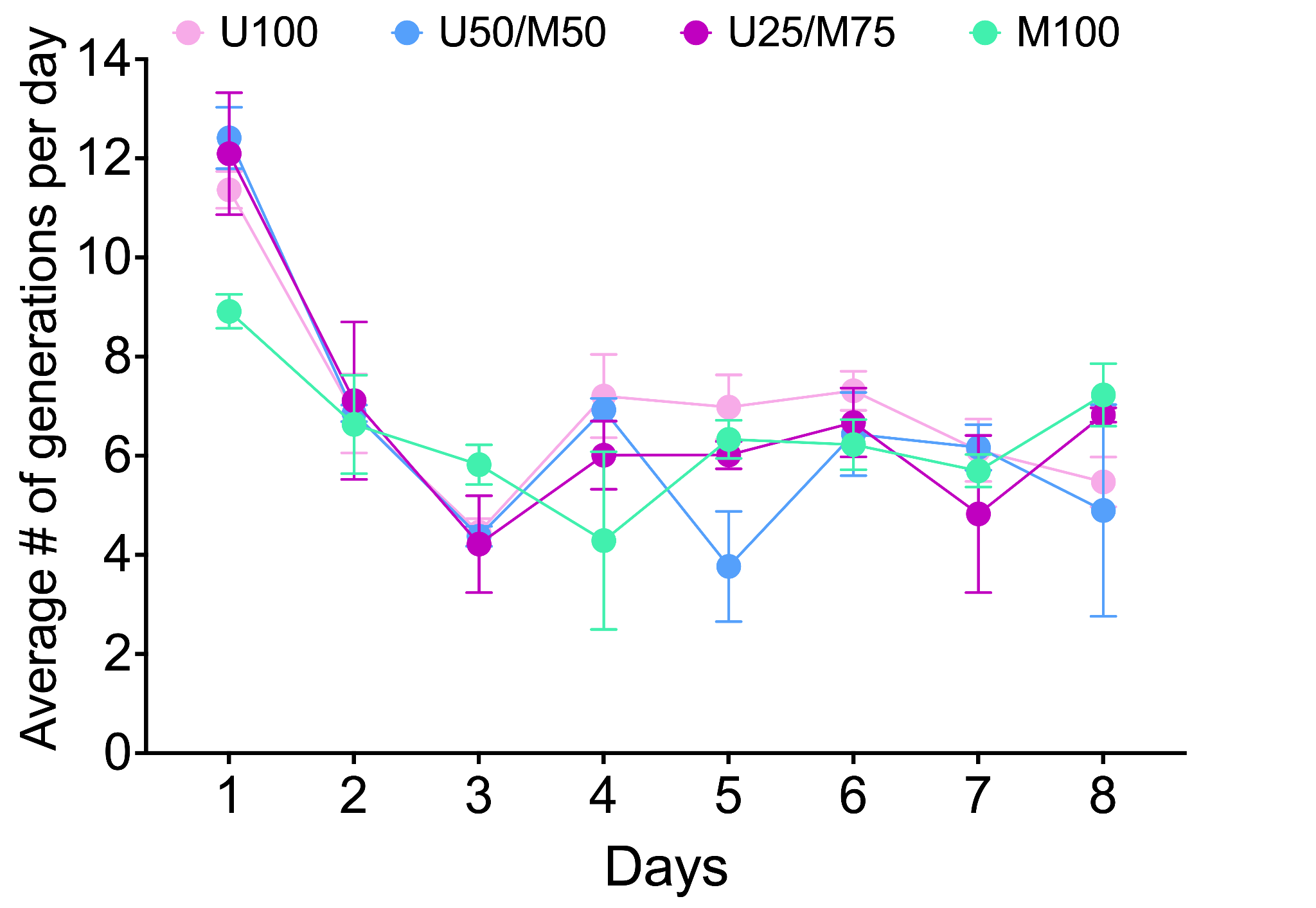

**Supplementary Figure 2. Number of generations across each life cycle.** We followed three cultures per treatment across eight days. Points show daily means ± SD. Mean daily counts across treatments were approximately six during days 4–8. We did not detect a treatment difference in total generations per cycle (*F*_3,8_ = 2.50, *p* = 0.134, ANOVA).

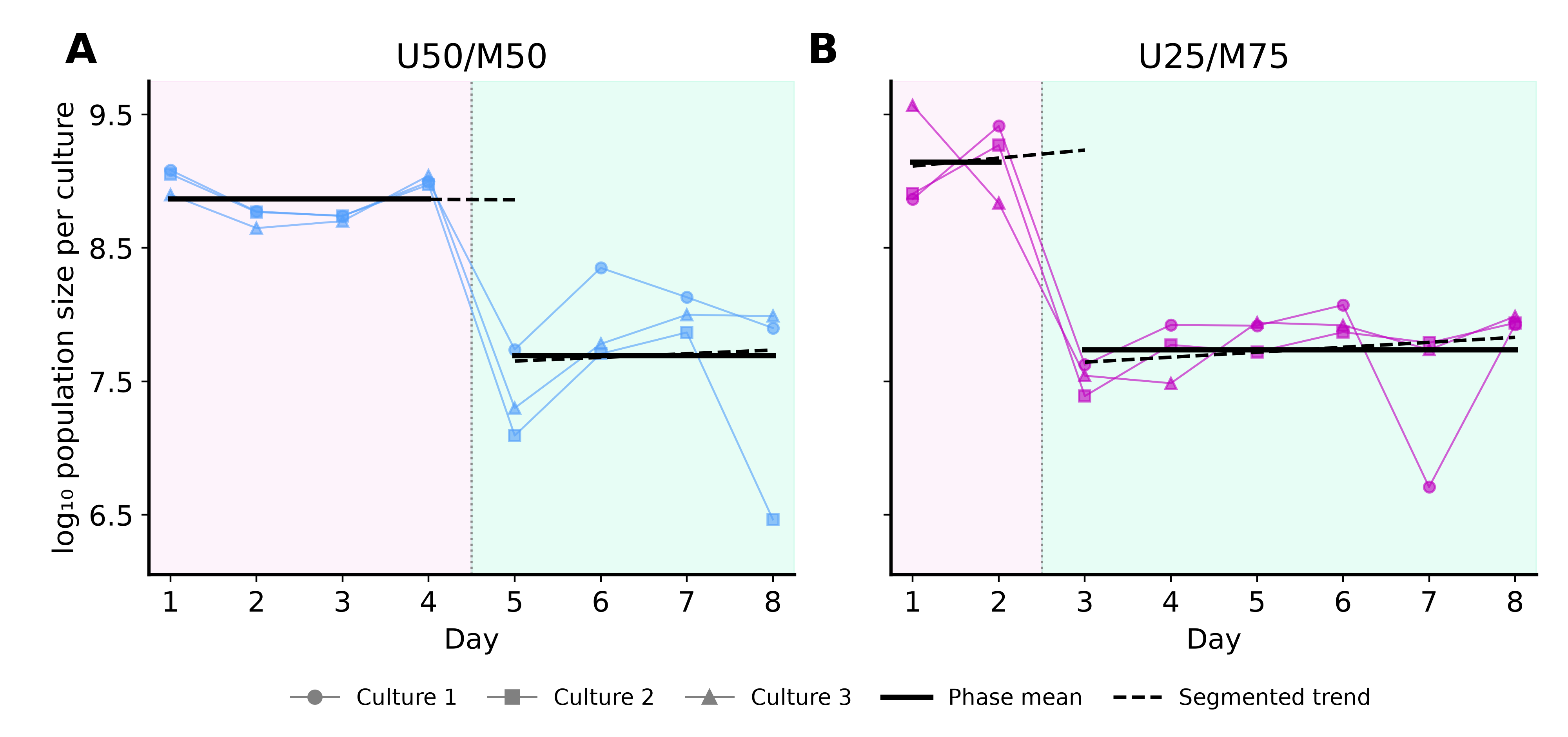

**Supplementary Figure 3. Culture-level population dynamics and sensitivity to within-phase trends.** Points and connecting lines show three cultures per treatment followed across eight days in U50/M50 (A) and U25/M75 (B). Pink and green shading mark unicellular and multicellular phases. Solid black lines show mean fitted phase levels from culture-specific piecewise-constant regressions. Paired comparisons of culture phase means gave Holm-adjusted p = 0.0333 for U50/M50 and p = 0.00108 for U25/M75 (n = 3 cultures each; adjustment includes the U100/M100 comparison in Figure 2). Dashed black lines show mean segmented fits allowing separate slopes within each phase, with pre-switch trends extrapolated to the first multicellular measurement (day 5 or day 3). For this sensitivity analysis, the mean pre-minus-post line separation was 1.208 log_10_ units for U50/M50 (95% CI, 0.345–2.072; t = 6.02, df = 2; unadjusted p = 0.0265) and 1.589 for U25/M75 (95% CI, −0.654–3.833; t = 3.05, df = 2; unadjusted p = 0.0929). All tests were two-sided; sensitivity intervals are pointwise and unadjusted. Thus, phase-average population sizes were lower during multicellular growth, whereas the trend-adjusted separation for U25/M75 remained uncertain. Because only two measurements precede the U25/M75 switch, the segmented estimate has less precision than the U50/M50 comparison. It is, however, consistent with a rapid switch multicellularity upon β-estradiol induction.

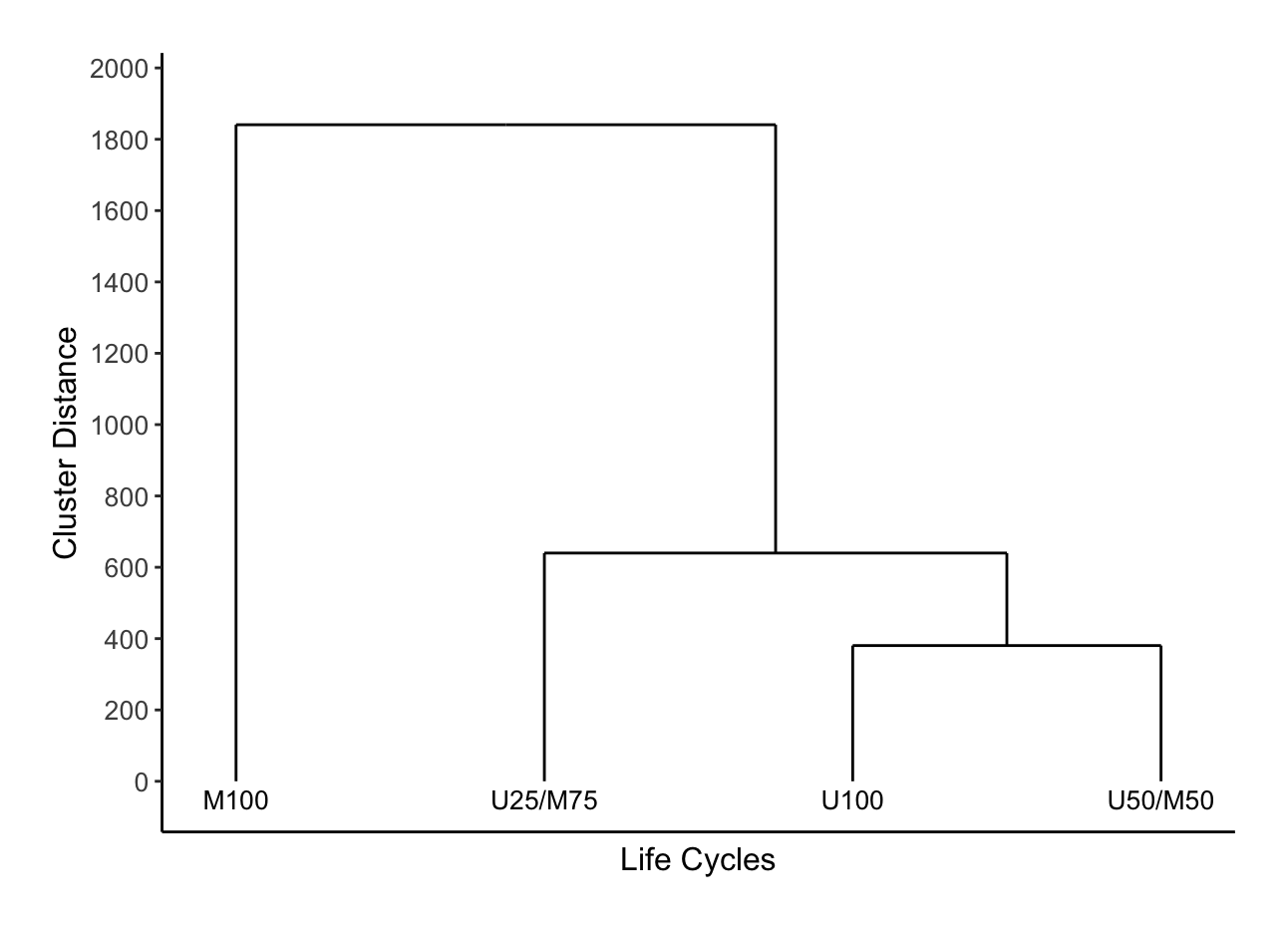

**Supplementary Figure 4. Facultative life cycles resemble obligately unicellular populations in mutant lineage survival, even when most of the life cycle is spent in a multicellular phase.** We represented each life cycle by 36 mean mutant-survival counts spanning nine growth effects and four group-level selection effects (Figure 5). Each mean averages three batches of 1,000 simulations and counts lineages surviving eight rounds of growth and transfer. We used agglomerative hierarchical clustering with Euclidean distances and Ward's minimum-variance linkage, without standardizing the features. Branch heights are Ward linkage distances on the mean-survival-count scale (out of 1,000). Supplementary Code 5 reproduces this analysis.
