## Supplementary Wright Fisher Model for "Obligate multicellularity circumvents population genetic barriers to collective-level adaptation": build_figure4.py

```
from pathlib import Path
import csv, json, hashlib
import numpy as np
import openpyxl
import matplotlib
matplotlib.use('Agg')
import matplotlib.pyplot as plt

import argparse
from make_figure4a import load_means

parser = argparse.ArgumentParser(description="Reproduce the five-panel Figure 4 from model trajectories and source data.")
parser.add_argument('--workbook', type=Path, default=Path('../Supplementary_Data.xlsx'))
parser.add_argument('--input', type=Path, default=Path('outputs/data/two_cycles_50pct.csv'))
parser.add_argument('--output-dir', type=Path, default=Path('outputs/figure4'))
args = parser.parse_args()
out = args.output_dir
out.mkdir(parents=True, exist_ok=True)
wb = openpyxl.load_workbook(args.workbook, data_only=True)
load_means(args.input)
order=['U100','U50/M50','U25/M75','M100']
colors={'U100':'#f7abe8','U50/M50':'#55a0fb','U25/M75':'#c000c0','M100':'#30dea5'}
rows=list(csv.DictReader(args.input.open()))
replicates={name:len([x for x in rows if x['life_cycle']==name and int(x['day'])==0]) for name in order}
means={name:np.array([np.mean([float(x['frequency']) for x in rows if x['life_cycle']==name and int(x['day'])==day]) for day in range(17)]) for name in order}
comp=[float(wb['figure4_ploidycompetition'][f'B{r}'].value) for r in range(2,11)]
reevolved={c:[float(wb['figure4_ploidy_reevolved'][f'{c}{r}'].value) for r in range(2,5)] for c in ['B','C','D']}
parent_labels=[str(wb['figure4_ploidy_reevolved'][f'{c}1'].value) for c in ['B','C','D']]
settling=[[float(wb['figure4_relaxedvssettling'].cell(3,c).value) for c in [2,3,4]], [float(wb['figure4_relaxedvssettling'].cell(3,c).value) for c in [5,6,7]]]
radii=[[float(wb['figure4_tetraploidy_be'].cell(2,c).value) for c in range(2,7)], [float(wb['figure4_tetraploidy_be'].cell(2,c).value) for c in range(7,12)]]
plt.rcParams.update({'font.family':'sans-serif','font.sans-serif':['Arial','DejaVu Sans'],'font.size':11,'axes.labelsize':13,'axes.linewidth':1,'xtick.labelsize':11,'ytick.labelsize':11,'pdf.fonttype':42,'svg.fonttype':'none'})

def clean(ax,label):
    ax.spines[['top','right']].set_visible(False)
    ax.tick_params(direction='out',length=3.5,width=1,pad=3)
    ax.text(-0.17 if label!='A' else -0.085,1.025,label,transform=ax.transAxes,fontsize=16,fontweight='bold',va='bottom',ha='left')

def simulation(ax,label,two=True):
    end=16 if two else 8
    ax.axhline(.5,color='black',ls='--',lw=1,zorder=0)
    if two:
        ax.axvline(8,color='#bcbcbc',lw=.9,ls=':',zorder=0)
        for x,s in [(4,'Cycle 1'),(12,'Cycle 2')]:ax.text(x,1.035,s,ha='center',va='bottom',fontsize=10,color='#666666')
    for name in order:ax.plot(np.arange(end+1),means[name][:end+1],color=colors[name],lw=2,label=name)
    ax.set(xlim=(0,end),ylim=(0,1.08),xlabel='Days',ylabel='Frequency of 4N')
    ax.set_xticks(np.arange(0,end+1,2));ax.set_yticks([0,.5,1],labels=['0.0','0.5','1.0'])
    ax.legend(loc='lower left',frameon=False,fontsize=10,ncol=2,handlelength=1.3,columnspacing=1.2)
    clean(ax,label)

def direct(ax,label):
    offsets=np.array([0,0,.09,-.09,-.27,0,0,.27,0])
    ax.scatter(offsets,comp,color='#505050',s=27,zorder=3)
    ax.hlines(np.mean(comp),-.26,.26,color='#505050',lw=1.5)
    ax.set(xlim=(-.85,.85),ylim=(0,2.5),ylabel='4N fitness relative to 2N (r)')
    ax.set_xticks([0],['U25/M75']);clean(ax,label)

def reev(ax,label):
    for i,c in enumerate(['B','C','D']):
        v=reevolved[c]
        ax.scatter(i+np.array([-.07,0,.07]),v,color='#505050',s=26,zorder=3)
        ax.hlines(np.mean(v),i-.26,i+.26,color='#505050',lw=1.5)
    ax.set(xlim=(-.65,2.65),ylim=(0,8),xlabel='Parent population',ylabel='DNA content')
    ax.set_xticks([0,1,2],parent_labels);ax.set_yticks([0,2,4,6,8],labels=['0N','2N','4N','6N','8N']);clean(ax,label)

def settle(ax,label):
    for i,v in enumerate(settling):
        ax.scatter(i+np.array([-.09,0,.09]),v,color='#2929ff',s=26,zorder=3)
        ax.hlines(np.mean(v),i-.26,i+.26,color='#2929ff',lw=1.1)
    ax.set(xlim=(-.65,1.65),ylim=(0,5),ylabel='DNA content',xlabel='Settling selection')
    ax.set_xticks([0,1],['Without','With']);ax.set_yticks(range(6),labels=[f'{i}N' for i in range(6)]);clean(ax,label)

def radius(ax,label):
    offsets=[-.08,-.16,.03,.16,.08]
    for i,v in enumerate(radii):
        ax.scatter(i+np.array(offsets),v,color='#9217ed',s=26,zorder=3)
        ax.hlines(np.mean(v),i-.26,i+.26,color='#9217ed',lw=1.5)
    ax.plot([0,0,1,1],[23,23.6,23.6,23],color='#555555',lw=.7)
    ax.text(.5,23.7,'****',ha='center',va='bottom',fontsize=12,fontweight='bold')
    ax.set(xlim=(-.65,1.65),ylim=(0,26),ylabel='Cluster radius (µm)',xlabel='β-estradiol')
    ax.set_xticks([0,1],['Without','With']);ax.set_yticks([0,5,10,15,20,25]);clean(ax,label)

fig=plt.figure(figsize=(10.6,6.95))
grid=fig.add_gridspec(2,3,left=.074,right=.985,bottom=.10,top=.935,wspace=.43,hspace=.58)
simulation(fig.add_subplot(grid[0,:2]),'A')
direct(fig.add_subplot(grid[0,2]),'B')
reev(fig.add_subplot(grid[1,0]),'C')
settle(fig.add_subplot(grid[1,1]),'D')
radius(fig.add_subplot(grid[1,2]),'E')
for ext in ['png','pdf','svg']:fig.savefig(out/f'Figure_4.{ext}',dpi=400)
plt.close(fig)

# Retain a separate model panel for editing and reproduction.
fig,ax=plt.subplots(figsize=(6.7,3.25));fig.subplots_adjust(left=.105,right=.98,bottom=.19,top=.85)
simulation(ax,'A')
for ext in ['png','pdf','svg']:fig.savefig(out/f'Figure_4_model_two_cycles.{ext}',dpi=400)
plt.close(fig)

with (out/'Figure_4_model_means.csv').open('w',newline='') as f:
    writer=csv.writer(f);writer.writerow(['life_cycle','day','mean_frequency','n_simulations'])
    for name in order:
        writer.writerows([[name,i,float(v),replicates[name]] for i,v in enumerate(means[name])])
record={'panel_mapping':{'A':'model','B':'direct competition','C':'re-evolution','D':'settling control','E':'induced unicellularity'},'experimental_values':{'B':comp,'C':reevolved,'D':settling,'E':radii},'day8':{n:float(means[n][8]) for n in order},'day16':{n:float(means[n][16]) for n in order},'trajectories_per_regime':replicates}
record['source_workbook_sha256']=hashlib.sha256(args.workbook.read_bytes()).hexdigest()
record['trajectory_sha256']=hashlib.sha256(args.input.read_bytes()).hexdigest()
(out/'Figure_4_metadata.json').write_text(json.dumps(record,indent=2))
print(json.dumps({'day8':record['day8'],'day16':record['day16']},indent=2))
```
