## Supplementary Wright Fisher Model for "Obligate multicellularity circumvents population genetic barriers to collective-level adaptation": make_figure4a.py

```
"""Plot the two eight-day cycles of the 50% tetraploid simulation as Figure 4A."""

import argparse
import csv
import hashlib
import json
from pathlib import Path

import numpy as np

from wright_fisher_life_cycles import COLORS, validate_frequency

ORDER = ("U100", "U50/M50", "U25/M75", "M100")


def load_means(path):
    records = {}
    with Path(path).open(newline="") as handle:
        reader = csv.DictReader(handle)
        required = {"life_cycle", "replicate", "day", "frequency"}
        if not required.issubset(reader.fieldnames or []):
            raise ValueError("Input must contain life_cycle, replicate, day and frequency")
        for row in reader:
            name, rep, day = row["life_cycle"], int(row["replicate"]), int(row["day"])
            if name not in ORDER or rep < 0 or day < 0:
                raise ValueError("Invalid life-cycle, replicate or day identifier")
            key = (name, rep, day)
            if key in records:
                raise ValueError(f"Duplicate observation: {key}")
            records[key] = validate_frequency(row["frequency"])
    if not records:
        raise ValueError("Input contains no observations")
    means, counts = {}, {}
    for name in ORDER:
        reps = sorted({rep for lc, rep, _ in records if lc == name})
        if not reps:
            raise ValueError(f"Missing life cycle: {name}")
        values = []
        for rep in reps:
            keys = [(name, rep, day) for day in range(17)]
            if any(key not in records for key in keys):
                raise ValueError(f"Incomplete days 0-16 for {name}, replicate {rep}")
            trajectory = [records[key] for key in keys]
            if trajectory[0] != 0.5:
                raise ValueError("Figure 4A requires a 50% starting frequency")
            values.append(trajectory)
        means[name] = np.mean(values, axis=0)
        counts[name] = len(reps)
    return means, counts


def draw_panel(ax, means):
    ax.axvline(8, color="#bcbcbc", linestyle=":", linewidth=0.9)
    for x, label in [(4, "Cycle 1"), (12, "Cycle 2")]:
        ax.text(x, 1.035, label, ha="center", va="bottom", color="#666666")
    ax.axhline(0.5, color="black", linestyle="--", linewidth=1.5, zorder=1)
    for name in ORDER:
        ax.plot(range(17), means[name], color=COLORS[name], linewidth=1.6,
                label=name, solid_capstyle="round", zorder=2)
    ax.set(xlim=(0, 16), ylim=(0, 1.1), xlabel="Days", ylabel="Frequency of 4N")
    ax.set_xticks(range(0, 17, 2))
    ax.set_yticks([0, 0.5, 1.0], labels=["0.0", "0.5", "1.0"])
    for spine in ("top", "right"):
        ax.spines[spine].set_visible(False)
    ax.tick_params(direction="out", length=3, width=0.8, pad=2)
    ax.legend(loc="lower left", frameon=False, fontsize=7,
              handlelength=0.7, handletextpad=0.4, labelspacing=0.45)


def main():
    parser = argparse.ArgumentParser(description=__doc__)
    parser.add_argument("--input", type=Path, default=Path("outputs/data/two_cycles_50pct.csv"))
    parser.add_argument("--output-dir", type=Path, default=Path("outputs/figure4a"))
    args = parser.parse_args()
    means, counts = load_means(args.input)

    import matplotlib
    matplotlib.use("Agg")
    import matplotlib.pyplot as plt

    plt.rcParams.update({"font.family": "sans-serif", "font.sans-serif": ["Arial", "DejaVu Sans"],
                         "font.size": 10, "pdf.fonttype": 42, "svg.fonttype": "none"})
    fig, ax = plt.subplots(figsize=(6.7, 3.25))
    fig.subplots_adjust(left=0.105, right=0.98, bottom=0.19, top=0.85)
    draw_panel(ax, means)
    args.output_dir.mkdir(parents=True, exist_ok=True)
    for suffix in ("png", "pdf", "svg"):
        fig.savefig(args.output_dir / f"Figure_4A.{suffix}", dpi=400)
    plt.close(fig)
    with (args.output_dir / "Figure_4A_means.csv").open("w", newline="") as handle:
        writer = csv.writer(handle)
        writer.writerow(["life_cycle", "day", "mean_frequency", "n_simulations"])
        for name in ORDER:
            writer.writerows([name, day, float(value), counts[name]]
                             for day, value in enumerate(means[name]))
    metadata = {"source_file": str(args.input),
                "source_sha256": hashlib.sha256(args.input.read_bytes()).hexdigest(),
                "replicates_per_regime": counts, "days": [0, 16], "starting_frequency": 0.5,
                "summary": "Arithmetic mean across simulations; no uncertainty band.",
                "colors": COLORS, "matplotlib": matplotlib.__version__}
    (args.output_dir / "Figure_4A_metadata.json").write_text(json.dumps(metadata, indent=2) + "\n")
    print(f"Wrote Figure 4A and its plotted means to {args.output_dir}")


if __name__ == "__main__":
    main()
```
