## Supplementary Wright Fisher Model for "Obligate multicellularity circumvents population genetic barriers to collective-level adaptation": README.md

# Wright-Fisher simulation and Figure 4

We simulate diploid-tetraploid competition during unicellular and multicellular growth. Each day combines a deterministic change in relative frequency with binomial transfer. Extinction and fixation are absorbing states; the model contains no mutation or migration.

## Run the model and plot the data

Tested with Python 3.12.14 on macOS arm64. From the extracted package directory:

```sh
python -m pip install -r requirements.txt
python wright_fisher_life_cycles.py --output-dir outputs
python build_figure4.py --workbook ../Supplementary_Data.xlsx --input outputs/data/two_cycles_50pct.csv --output-dir outputs/figure4
```

The simulation does not require the workbook. The composite plotting script reads the accompanying workbook for experimental panels B-E and generates a plot from the Figure 4 data. It also exports the model panel separately and writes all 68 plotted model means. The workbook is read without modification. Plot appearance depends on available fonts; the manuscript figure uses Arial.

For a model-only plot without the experimental panels or workbook:

```sh
python make_figure4a.py --input outputs/data/two_cycles_50pct.csv --output-dir outputs/figure4a
```

This compact model-only plot uses the same trajectories and arithmetic means. Use `build_figure4.py` to plot the model and experimental panels together.

## Fitness inputs

Default daily log-odds increments favoring tetraploids are:

| Context | Mean r per day | Sample SD | n |
| --- | ---: | ---: | ---: |
| Unicellular, reference-based contrasts | -0.07377261129458766 | 0.363678235015511 | 3 |
| Multicellular, published mixotrophic competitions with settling | 0.4712857404482067 | 0.10505681185297748 | 4 |

The `figure4_fitness_inputs` worksheet supplies these means. The 2N and 4N assays used the same GFP competitor. Unicellular values are calculated from the counts in `figure4A_unicellular_counts`: four GFP fractions are averaged per endpoint, and the change in reference log odds is contrasted between test ploidies. The three contrasts pair replicate indices and share day-0 baselines.

Multicellular rates come from Tong et al., *Genome duplication in a long-term multicellularity evolution experiment*, Nature 639, 691-699 (2025), Figure 3g, DOI [10.1038/s41586-025-08689-6](https://doi.org/10.1038/s41586-025-08689-6). We use the four `Condition = PM`, `Selection = S` rows in `source/Fig3g.csv`. The data were retrieved from [WGD_in_MuLTEE](https://github.com/ktong25/WGD_in_MuLTEE), commit `aa090bbd9163dda490a5ede716bdd063270e9cd6`, path `data/source_data/Fig3g.csv`.

For tetraploid volume fractions p0 and p3, the published daily rate is `[logit(p3) - logit(p0)] / 3`. It already includes growth and settling. The published daily transfer settled 1.5 mL for 3 minutes and transferred the bottom 50 µL into fresh medium. This differs from the model's idealized 1/32 transfer. The sources use distinct assays and are not paired or pooled.

The four published mixotrophic competitions without settling have mean r = -0.04959075789050517 per day and SD = 0.16144187117095254. They are used only for the comparison in the Results; their endpoints and calculations are in `figure4_fitness_inputs`.

`input_estimates.json` records the input values and workbook checksum. The model treats the means as fixed and does not propagate assay uncertainty.

## Model and schedules

The population contains 100 million cells before transfer. Multicellular sampling treats these as 10 million clonal groups of 10 cells. Transfer retains 1/32 of the units: 3,125,000 cells on a unicellular day or 312,500 groups on a multicellular day.

For an interior tetraploid frequency p, selection gives `logit(p_selected) = logit(p) + r`. Transfer then samples `Binomial(n, p_selected) / n`. Regrowth adds no frequency change beyond the daily rate already applied. Rates remain constant within each growth phase; relaxed selection on transition days is not represented explicitly.

| Treatment | Eight-day schedule |
| --- | --- |
| U100 | U U U U U U U U |
| M100 | M M M M M M M M |
| U25/M75 | U U M M M M M M |
| U50/M50 | U U U U M M M M |

Figure 4A repeats each schedule twice. It shows arithmetic means across 1,000 trajectories per regime on days 0-16, initially 50% tetraploid, without an uncertainty band. The horizontal dashed line marks the starting frequency; the vertical dotted line separates the cycles.

## Outputs and reuse

Paths are relative to the chosen output directory.

| Output | Content |
| --- | --- |
| `data/two_cycles_50pct.csv` | Days 0-16, initially 50% tetraploid; 1,000 trajectories per regime by default. |
| `data/two_cycles_1pct.csv` | Same design, initially 1% tetraploid; a diagnostic analysis. |
| `data/extinction_vs_start_cells.csv` | Extinction at day 8 for 50 rounded logarithmically spaced starting counts from 10 to 10,000 cells, 1,000 trials per count and regime; a diagnostic analysis. |
| `plots/` | Extinction curve and two diagnostic log-frequency plots. Zeros remain zero in CSVs and are masked on log axes. |
| `run_metadata.json` | Parameters, seeds, software versions, runtime and data checksums. |
| `figure4/Figure_4.*` | Composite plot of the Figure 4 data. |
| `figure4/Figure_4_model_two_cycles.*` | Model trajectory panel. |
| `figure4/Figure_4_model_means.csv` | All 68 plotted model means and replicate counts. |
| `figure4/Figure_4_metadata.json` | Experimental values, panel mapping, source checksums and model endpoints. |

Use `--r-uni` and `--r-multi` to override either finite daily log-odds increment. The simulation also accepts `--two-cycles-reps`, `--extinction-points`, `--extinction-reps` and `--seed-offset`. `--skip-extinction` omits the extinction grid; `--skip-two-cycles` omits both trajectory analyses. Use a separate output directory for each parameter set. The plotting scripts require a 50% starting frequency and complete days 0-16 for each trajectory. They are intended for the four supplied regimes.

NumPy PCG64 generates the random draws. Trajectory replicate i uses seed `9000 + seed_offset + i`. Extinction replicate i at starting count c uses `5000 + 1000000*c + seed_offset + i`. Indices start at zero. Seeds are shared across regimes and trajectory starting frequencies, so comparisons do not use independent random streams. The default offset is zero; the pinned dependencies reproduce the reference outputs.
