## Supplementary Wright Fisher Model for "Obligate multicellularity circumvents population genetic barriers to collective-level adaptation": wright_fisher_life_cycles.py

```
"""Daily selection and binomial transfer of diploid and tetraploid lineages."""

import argparse
import csv
from dataclasses import asdict, dataclass
import hashlib
import json
import math
from pathlib import Path
import platform
import time

import numpy as np


N_CELLS = 10**8
GROUP_SIZE = 10
N_GROUPS = N_CELLS // GROUP_SIZE
DILUTION = 32
# Mean daily selection rates are listed in input_estimates.json.
R_UNI_FROM_COUNTS = -0.07377261129458766
R_MULTI_FROM_TONG = 0.4712857404482067
SCHEDULES = {
    "U100": tuple("UUUUUUUU"),
    "M100": tuple("MMMMMMMM"),
    "U25/M75": tuple("UUMMMMMM"),
    "U50/M50": tuple("UUUUMMMM"),
}
COLORS = {
    "U100": "#f7abe8",
    "U50/M50": "#55a0fb",
    "U25/M75": "#c000c0",
    "M100": "#41f0ae",
}


@dataclass(frozen=True)
class PhaseParams:
    """Tetraploid-minus-diploid log-odds increments per day."""

    r_uni: float = R_UNI_FROM_COUNTS
    r_multi: float = R_MULTI_FROM_TONG

    def __post_init__(self):
        if not all(math.isfinite(r) for r in (self.r_uni, self.r_multi)):
            raise ValueError("Selection rates must be finite")


def validate_frequency(p):
    p = float(p)
    if not math.isfinite(p) or not 0 <= p <= 1:
        raise ValueError("Frequency must be finite and between zero and one")
    return p


def apply_selection(p, r):
    """Apply logit(p_next) = logit(p) + r, preserving absorbing endpoints."""
    p = validate_frequency(p)
    if not math.isfinite(r):
        raise ValueError("Selection rate must be finite")
    if p == 0 or p == 1:
        return p
    logit_prime = math.log(p / (1 - p)) + r
    # Avoid overflow in the far tail.
    if logit_prime < -700:
        odds = math.exp(logit_prime)
        return odds / (1 + odds)
    return 1.0 / (1.0 + math.exp(-logit_prime))


def get_rng(seed=42):
    return np.random.default_rng(seed)


def passage_with_drift(p, units_total, dilution, rng):
    """Transfer a binomial sample of cells or clonal groups."""
    p = validate_frequency(p)
    if (not isinstance(units_total, (int, np.integer)) or
            not isinstance(dilution, (int, np.integer)) or
            units_total <= 0 or dilution <= 0):
        raise ValueError("Population and dilution must be positive integers")
    n = units_total // dilution
    if n <= 0:
        raise ValueError("Passaged sample size must be positive")
    if p == 0 or p == 1:
        return p
    return int(rng.binomial(n, p)) / n


def simulate_schedule(phases, p0, params, seed=42):
    """Return tetraploid frequencies before growth and after each daily transfer."""
    phases = tuple(phase.upper() for phase in phases)
    if any(phase not in ("U", "M") for phase in phases):
        raise ValueError("Each phase must be U or M")
    p = validate_frequency(p0)
    rng = get_rng(seed)
    freqs = [p]
    for phase in phases:
        r = params.r_uni if phase == "U" else params.r_multi
        units = N_CELLS if phase == "U" else N_GROUPS
        p = passage_with_drift(apply_selection(p, r), units, DILUTION, rng)
        freqs.append(p)
    return freqs


def write_csv(path, header, rows):
    with path.open("w", newline="") as handle:
        writer = csv.writer(handle)
        writer.writerow(header)
        writer.writerows(rows)


def run_trajectories(params, p0, reps, seed_offset):
    rows = []
    means = {}
    for name, schedule in SCHEDULES.items():
        trajectories = []
        for i in range(reps):
            freqs = simulate_schedule(schedule * 2, p0, params,
                                      seed=9000 + seed_offset + i)
            trajectories.append(freqs)
            rows.extend([name, i, day, value] for day, value in enumerate(freqs))
        means[name] = np.mean(trajectories, axis=0)
    return rows, means


def run_extinction_grid(params, points, reps, seed_offset):
    starts = np.unique(np.round(np.logspace(1, 4, points)).astype(int))
    rows = []
    fractions = {}
    for name, schedule in SCHEDULES.items():
        fractions[name] = []
        for cells in starts:
            extinct = 0
            for i in range(reps):
                seed = 5000 + 1000000 * int(cells) + seed_offset + i
                freqs = simulate_schedule(schedule, cells / N_CELLS, params, seed)
                lost = int(freqs[-1] == 0)
                extinct += lost
                rows.append([name, int(cells), i, lost])
            fractions[name].append(extinct / reps)
    return rows, starts, fractions


def positive_integer(value):
    n = int(value)
    if n < 1:
        raise argparse.ArgumentTypeError("Value must be a positive integer")
    return n


def main():
    parser = argparse.ArgumentParser(description=__doc__)
    parser.add_argument("--r-uni", type=float, default=R_UNI_FROM_COUNTS,
                        help="Daily increment; default is the unrounded source-count mean")
    parser.add_argument("--r-multi", type=float, default=R_MULTI_FROM_TONG,
                        help="Daily increment; default is the Tong et al. mixotrophic settling mean")
    parser.add_argument("--seed-offset", type=int, default=0)
    parser.add_argument("--extinction-points", type=positive_integer, default=50)
    parser.add_argument("--extinction-reps", type=positive_integer, default=1000)
    parser.add_argument("--two-cycles-reps", type=positive_integer, default=1000)
    parser.add_argument("--skip-extinction", action="store_true")
    parser.add_argument("--skip-two-cycles", action="store_true")
    parser.add_argument("--output-dir", type=Path, default=Path("outputs"))
    args = parser.parse_args()
    if args.seed_offset < 0:
        parser.error("--seed-offset must be nonnegative")
    if args.skip_extinction and args.skip_two_cycles:
        parser.error("At least one analysis must be selected")
    try:
        params = PhaseParams(args.r_uni, args.r_multi)
    except ValueError as error:
        parser.error(str(error))

    import matplotlib
    matplotlib.use("Agg")
    import matplotlib.pyplot as plt

    started = time.perf_counter()
    data_dir, plot_dir = args.output_dir / "data", args.output_dir / "plots"
    data_dir.mkdir(parents=True, exist_ok=True)
    plot_dir.mkdir(parents=True, exist_ok=True)
    written = []
    if not args.skip_extinction:
        rows, starts, fractions = run_extinction_grid(
            params, args.extinction_points, args.extinction_reps, args.seed_offset)
        path = data_dir / "extinction_vs_start_cells.csv"
        write_csv(path, ["life_cycle", "start_cells", "replicate", "extinct_end"], rows)
        written.append(path)
        fig, ax = plt.subplots(figsize=(5, 5))
        for name in SCHEDULES:
            ax.plot(starts, fractions[name], "o-", color=COLORS[name], label=name)
        ax.set(xscale="log", ylim=(0, 1.02), xlabel="Starting tetraploid cells",
               ylabel="Extinction fraction by day 8")
        ax.legend(frameon=False)
        fig.tight_layout()
        fig.savefig(plot_dir / "extinction_vs_start_cells.png", dpi=200)
        plt.close(fig)

    if not args.skip_two_cycles:
        for p0, tag in [(0.5, "50pct"), (0.01, "1pct")]:
            rows, means = run_trajectories(params, p0, args.two_cycles_reps, args.seed_offset)
            path = data_dir / f"two_cycles_{tag}.csv"
            write_csv(path, ["life_cycle", "replicate", "day", "frequency"], rows)
            written.append(path)
            fig, ax = plt.subplots(figsize=(5, 5))
            for name, mean in means.items():
                # Zero means are undefined on a log axis; do not alter the data.
                ax.plot(range(17), np.ma.masked_less_equal(mean, 0),
                        color=COLORS[name], label=name)
            ax.set(yscale="log", ylim=(1e-9, 1.02), xlim=(0, 16),
                   xlabel="Days", ylabel="Tetraploid frequency")
            ax.legend(frameon=False)
            fig.tight_layout()
            fig.savefig(plot_dir / f"two_cycles_{tag}_log.png", dpi=200)
            plt.close(fig)

    metadata = {
        "parameters": asdict(params), "carrying_capacity_cells": N_CELLS,
        "group_size_cells": GROUP_SIZE, "dilution": DILUTION,
        "schedules": {name: "".join(schedule) for name, schedule in SCHEDULES.items()},
        "absorbing_endpoints": True, "seed_offset": args.seed_offset,
        "trajectory_seed": "9000 + seed_offset + replicate",
        "extinction_seed": "5000 + 1000000 * start_cells + seed_offset + replicate",
        "seeds_shared_across_regimes": True,
        "two_cycles_replicates": None if args.skip_two_cycles else args.two_cycles_reps,
        "extinction_replicates": None if args.skip_extinction else args.extinction_reps,
        "extinction_requested_points": None if args.skip_extinction else args.extinction_points,
        "fitness_input_description": "Defaults are the unicellular estimate from this study and the multicellular settling estimate from Tong et al. (2025); command-line rates override them.",
        "python": platform.python_version(), "numpy": np.__version__,
        "matplotlib": matplotlib.__version__, "rng": "numpy.random.PCG64",
        "script_sha256": hashlib.sha256(Path(__file__).read_bytes()).hexdigest(),
        "data_sha256": {p.name: hashlib.sha256(p.read_bytes()).hexdigest() for p in written},
        "runtime_seconds": time.perf_counter() - started,
    }
    (args.output_dir / "run_metadata.json").write_text(json.dumps(metadata, indent=2) + "\n")
    print(f"Wrote {len(written)} data files and diagnostic plots to {args.output_dir}")


if __name__ == "__main__":
    main()
```
