## Supplementary Mathematical Model for "Obligate multicellularity circumvents population genetic barriers to collective-level adaptation": as_regime.py

```
"""Eight-round lineage-survival model (Python implementation of ASregime.m).

Sampling branches, discrete growth times and partial-group handling follow
the model. Replicates are advanced together for speed; NumPy's
random stream is not intended to reproduce MATLAB's individual draws.
"""

from collections import Counter
from dataclasses import dataclass

import numpy as np


def matlab_round(values):
    """Round halfway cases away from zero, including negative normal draws."""
    values = np.asarray(values)
    return (np.sign(values) * np.floor(np.abs(values) + 0.5)).astype(np.int64)


class GrowthGrid:
    """Find the last grid point before total abundance exceeds f*N.

Binary search evaluates the same monotone threshold as the full MATLAB
grid search. The model uses a discrete growth-time grid.
    """

    def __init__(self, population=1_000_000, expansion=32, growth=1.0):
        if (population <= 0 or int(population) != population
                or expansion <= 1 or int(expansion) != expansion
                or not np.isfinite(growth) or growth <= 0):
            raise ValueError('Require positive integer N, integer f > 1 and finite growth > 0.')
        self.population = int(population)
        self.expansion = int(expansion)
        self.growth = float(growth)
        resident_time = np.log(float(expansion))
        mutant_time = resident_time / growth
        if growth == 1:
            start, stop, steps = .95*resident_time, 1.05*resident_time, 10
            step = .01*resident_time
        else:
            start, stop, steps = min(resident_time, mutant_time), max(resident_time, mutant_time), 10000
            step = (stop-start)/steps
        # Construct the specified uniform grid from both endpoints to limit
        # accumulated roundoff. Native runtimes may differ at the last bit.
        indices = np.arange(steps+1)
        self.times = np.where(indices <= steps//2,
                              start+indices*step, stop-(steps-indices)*step)
        self.times[steps//2] = (start+stop)/2
        self.resident_factor = np.exp(self.times)
        self.mutant_factor = np.exp(growth*self.times)

    def grow(self, mutants):
        mutants = np.asarray(mutants, dtype=np.int64)
        if np.any((mutants < 0) | (mutants > self.population)):
            raise ValueError('Growth requires a mutant count between zero and N.')
        target = self.expansion*self.population
        lo = np.zeros(mutants.shape, dtype=np.int64)
        hi = np.full(mutants.shape, len(self.times), dtype=np.int64)
        while np.any(lo < hi):
            mid = (lo+hi)//2
            safe = np.minimum(mid, len(self.times)-1)
            size = (mutants*self.mutant_factor[safe]
                    + (self.population-mutants)*self.resident_factor[safe])
            above = size > target
            active = lo < hi
            hi = np.where(active & above, mid, hi)
            lo = np.where(active & ~above, mid+1, lo)
        # Fail explicitly where MATLAB's find returns an empty index. Do not
        # silently substitute a continuous root or change the growth rule.
        if np.any(lo == len(self.times)):
            raise ArithmeticError('No growth-grid point exceeds capacity (undefined MATLAB index).')
        if np.any(lo == 0):
            raise ArithmeticError('First growth-grid point already exceeds capacity.')
        index = lo-1
        result = matlab_round(mutants*self.mutant_factor[index])
        return result, int(np.count_nonzero(lo == len(self.times)))


def sample_unweighted(total, successes, draws, rng, diagnostics=None, label='cells'):
    """Retain the engine's normal/hypergeometric threshold and variance."""
    successes = np.asarray(successes, dtype=np.int64)
    mean = draws*successes.astype(float)/total
    variance = (draws*successes.astype(float)/total * (1-successes/total)
                * (total-draws)/(total-1))
    normal = ((mean >= 10) & (draws-mean >= 10)
              & (draws/total <= .1) & (variance >= 9))
    output = np.empty(successes.shape, dtype=np.int64)
    output[normal] = matlab_round(rng.standard_normal(np.count_nonzero(normal))
                                 * np.sqrt(variance[normal])+mean[normal])
    other = ~normal
    output[other] = rng.hypergeometric(successes[other], total-successes[other], draws)
    if diagnostics is not None:
        diagnostics[label+'_normal'] += int(np.count_nonzero(normal))
        diagnostics[label+'_hypergeometric'] += int(np.count_nonzero(other))
        diagnostics['negative_normal_draws'] += int(np.count_nonzero(output < 0))
        diagnostics['draws_above_sample_size'] += int(np.count_nonzero(output > draws))
    return output


def sample_weighted(total, successes, draws, weight, rng, diagnostics=None):
    """Retain the weighted Poisson, binomial and high-frequency branches."""
    successes = np.asarray(successes, dtype=np.int64)
    probability = weight*successes/(weight*successes+total-successes)
    expected = draws*probability
    poisson = (expected < 10) & (draws/total < .1)
    binomial = (expected > 10) & (draws/total < .1) & (expected < draws-10)
    saturation = ~(poisson | binomial) & (expected/draws > .95)
    if np.any(~(poisson | binomial | saturation)):
        raise NotImplementedError('Weighted hypergeometric branch is not implemented in ASregime.m.')
    output = np.empty(successes.shape, dtype=np.int64)
    output[poisson] = rng.poisson(expected[poisson])
    output[binomial] = rng.binomial(draws, probability[binomial])
    output[saturation] = draws
    if diagnostics is not None:
        for name, mask in [('groups_poisson', poisson), ('groups_binomial', binomial),
                           ('groups_saturation', saturation)]:
            diagnostics[name] += int(np.count_nonzero(mask))
    return output


def retain_groups(mutants, group_size, selected):
    """Lose the partial group when fewer groups survive; otherwise keep cells.

The approximate weighted samplers can return more groups than exist. As in
MATLAB, this case retains the pre-selection mutant cell count.
    """
    groups = (mutants+group_size-1)//group_size
    return np.where(selected < groups, selected*group_size, mutants)


@dataclass
class SimulationResult:
    survival_count: int
    final_counts: np.ndarray
    diagnostics: dict


def simulate(regime, growth, weight, rng, runs=1000, population=1_000_000,
             expansion=32, group_size=20, initial_mutants=1):
    """Return survival after the supplied U=1 / M=2 schedule."""
    if not regime or any(phase not in (1, 2) for phase in regime):
        raise ValueError('Regime must be a nonempty sequence of 1s and 2s.')
    if (not np.isfinite(weight) or weight <= 0 or runs < 1 or int(runs) != runs
            or group_size < 1 or int(group_size) != group_size
            or initial_mutants < 0 or int(initial_mutants) != initial_mutants
            or initial_mutants > population):
        raise ValueError('Invalid weight, replicate count, group size or initial count.')
    grid = GrowthGrid(population, expansion, growth)
    population, expansion, group_size = int(population), int(expansion), int(group_size)
    mutants = np.full(int(runs), initial_mutants, dtype=np.int64)
    current_size = group_size
    diagnostics = Counter()
    for phase in regime:
        active = mutants > 0
        if not np.any(active):
            break
        grown, endpoint_uses = grid.grow(mutants[active])
        diagnostics['growth_no_crossing_endpoint'] += endpoint_uses
        diagnostics['growth_events'] += int(np.count_nonzero(active))
        if phase == 1:
            selected = sample_unweighted(expansion*population, grown, population,
                                         rng, diagnostics, 'cells')
            current_size = 1
        else:
            current_size = min(current_size*expansion, group_size)
            if population % current_size:
                raise ValueError('Number of selected groups must be an integer.')
            groups = (grown+current_size-1)//current_size
            total = expansion*population//current_size
            draws = population//current_size
            if weight == 1:
                selected_groups = sample_unweighted(total, groups, draws, rng, diagnostics, 'groups')
            else:
                selected_groups = sample_weighted(total, groups, draws, weight, rng, diagnostics)
            diagnostics['selected_groups_exceed_available'] += int(np.count_nonzero(selected_groups > groups))
            selected = retain_groups(grown, current_size, selected_groups)
        mutants[active] = selected
        diagnostics['post_selection_above_N'] += int(np.count_nonzero(selected > population))
        if np.any(selected > population):
            raise ArithmeticError('Sampling approximation produced more than N mutant cells.')
    return SimulationResult(int(np.count_nonzero(mutants > 0)), mutants, dict(diagnostics))
```
