## Supplementary Mathematical Model for "Obligate multicellularity circumvents population genetic barriers to collective-level adaptation": ASregime.m

```
function CountSurvival=ASregime(SelectionRegime,f,N,EffectGrowth,EffectSelection)
% SelectionRegime is a vector with entries of either 1 (uni) or 2 (multi) 
% f is the factor by which the population expands, f=32 is common
% N is the initial population size, N=10^6 is common
% EffectGrowth is the factor by which growth is changed
% EffectSelection is the factor by which mutant groups are more likely to
% be selected--- acting as if there are more of them
% Output:
% CountSurvival: the number of runs (out of 1000) that the mutant survives

% Parameters for simulations
NumberRuns=1000;
GroupSize=20;
MaxNumberRounds=length(SelectionRegime);

% Data storage
MatrixSelection=zeros(NumberRuns,MaxNumberRounds); % mutant number after selection
MatrixGrowth=zeros(NumberRuns,MaxNumberRounds); % mutant number after selection


CountSurvival=0;
for i1=1:NumberRuns
    % Display status
    %['Progress :', num2str(i1/NumberRuns),'%']
    
    % Initialize each run of the experiment
    NumberRounds=0;
    NumberMutants=1; % Begins with a single mutant
    NumberInitialGroups=N; % assumes that the first M round has groups of size 
    CurrentGroupSize=GroupSize;

    % Run the experiment
    while NumberMutants>0 & NumberRounds<MaxNumberRounds
        NumberRounds=NumberRounds+1;
        
        % Growth (either as U or M) so total cell number reaches f*N
        T1=log(f); % time for resident to reach f*N when alone
        T2=log(f)/EffectGrowth; % time for mutant to reach f*N when alone
        if T1<T2
            % mutant grows slower
            Trange=[T1:(T2-T1)/10000:T2];
        elseif T1>T2
            % mutant grows faster
            Trange=[T2:(T1-T2)/10000:T1];
        else
            % No growth effect, T1=T2
            Trange=[.95*T1:.01*T1:1.05*T1];
        end
        PopulationSize=NumberMutants*exp(EffectGrowth*Trange)+(N-NumberMutants)*exp(Trange);
        Index=find(PopulationSize>f*N,1); % find when population first reaches f*N
        Index=Index-1; % stop right before population reaches f*N
        Time=Trange(Index); % find time when it happens
        NumberMutants=round(NumberMutants*exp(EffectGrowth*Time));
        MatrixGrowth(i1,NumberRounds)=NumberMutants;


        if SelectionRegime(NumberRounds)==1
            % Selection as U
            % check if normal approximation works
            NormalApproximationWorks=true;
            if N*NumberMutants/(f*N)<10 || N-N*NumberMutants/(f*N)<10
                % expected number of mutants (and non-mutants) picked should be greater than 10
                NormalApproximationWorks=false;
            end
            if N/(f*N)>.1
                % fraction selected should be < 10%
                NormalApproximationWorks=false;
            end
            if N*NumberMutants/(f*N)*(1-NumberMutants/(f*N))*(f*N-N)/(f*N-1)<9
                %variance must be greater than 9
                NormalApproximationWorks=false;
            end
            %NormalApproximationWorks=false;
            if NormalApproximationWorks
                Mean=N*NumberMutants/(f*N);
                Std=sqrt(N*NumberMutants/(f*N)*(1-NumberMutants/(f*N))*(f*N-N)/(f*N-1));
                NumberMutants = round(randn(1,1)*Std+Mean);
            else
                NumberMutants = hygernd(f*N,NumberMutants,N);
            end
            MatrixSelection(i1,NumberRounds)=NumberMutants;
            CurrentGroupSize=1; % used if later selection regime is multi

        elseif SelectionRegime(NumberRounds)==2
            % Growth as M, convert number of cells into number of groups
            if CurrentGroupSize*f<GroupSize
                % groups did not have time to reach typical group size
                CurrentGroupSize=CurrentGroupSize*f;
            else
                % assume the life cycles are realized in the population
                CurrentGroupSize=GroupSize;
            end
            NumberMutantGroups=ceil(NumberMutants/CurrentGroupSize);

            % Selection as M
            NumberTotalGroups=floor(f*N/CurrentGroupSize);
            NumberGroupsToSelect=N/CurrentGroupSize;
            if EffectSelection ~= 1
                % check what approximation works
                ProbabilitySelectMutant=EffectSelection * NumberMutantGroups/(EffectSelection * NumberMutantGroups+NumberTotalGroups-NumberMutantGroups);
                Lambda=NumberGroupsToSelect* ProbabilitySelectMutant; % expected number of mutant groups selected
                % Decide which approximation to use
                if Lambda<10 && NumberGroupsToSelect/NumberTotalGroups<.1
                    % use Poisson approximation
                    NewNumberMutantGroups =poissrnd(Lambda);
                elseif Lambda>10 && NumberGroupsToSelect/NumberTotalGroups<.1 && Lambda<NumberGroupsToSelect-10
                    % use Binomial approximation
                    NewNumberMutantGroups =binornd(NumberGroupsToSelect,ProbabilitySelectMutant);
                else
                    if Lambda/NumberGroupsToSelect>.95
                        % expected number of mutant groups selected is close to 1
                        NewNumberMutantGroups=NumberGroupsToSelect;
                    else
                        error('Need to implement the weighted hypergeometric distribution directly');
                    end
                end
            else
                % All groups are equally likely to be picked
                % check if normal approximation works
                NormalApproximationWorks=true;
                if NumberGroupsToSelect*NumberMutantGroups/NumberTotalGroups<10 || NumberGroupsToSelect-NumberGroupsToSelect*NumberMutantGroups/NumberTotalGroups<10
                    % expected number of mutants (and non-mutants) picked should be greater than 10
                    NormalApproximationWorks=false;
                end
                if NumberGroupsToSelect/NumberTotalGroups>.1
                    % fraction selected should be < 10%
                    NormalApproximationWorks=false;
                end
                if NumberGroupsToSelect*NumberMutantGroups/NumberTotalGroups*(1-NumberMutantGroups/NumberTotalGroups)*(NumberTotalGroups-NumberGroupsToSelect)/(NumberTotalGroups-1)<9
                    %variance must be greater than 9
                    NormalApproximationWorks=false;
                end
                if NormalApproximationWorks
                    Mean=NumberGroupsToSelect*NumberMutantGroups/NumberTotalGroups;
                    Std=sqrt(NumberGroupsToSelect*NumberMutantGroups/NumberTotalGroups*(1-NumberMutantGroups/NumberTotalGroups)*(NumberTotalGroups-NumberGroupsToSelect)/(NumberTotalGroups-1));
                    NewNumberMutantGroups = round(randn(1,1)*Std+Mean);
                else
                    NewNumberMutantGroups = hygernd(NumberTotalGroups,NumberMutantGroups,NumberGroupsToSelect);
                end
            end
            if NewNumberMutantGroups<NumberMutantGroups
                % not all groups are selected, assumes hybrid group lost
                NumberMutants=NewNumberMutantGroups*CurrentGroupSize;
            end
            MatrixSelection(i1,NumberRounds)=NumberMutants;

        else
            % Selection regime was neither a 1 or 2
            error('Selection regime should only contain 1s or 2s as entries')
        end
    
    end

    if NumberMutants>0
        % the mutant is still present
        CountSurvival=CountSurvival+1;
    end
end
        
end
    
```
