## Supplementary Mathematical Model for "Obligate multicellularity circumvents population genetic barriers to collective-level adaptation": cycle_fitness.m

```
function [W, log2W] = cycle_fitness(regime, f, growth, weight)
% Relative cycle fitness in the deterministic, rare-mutant approximation.
% growth multiplies the resident exponential growth rate, not yield per round.
% Group-selection weight acts only in multicellular rounds (regime == 2).
% Discrete groups, stochastic loss and finite-frequency saturation are excluded.

validateattributes(regime, {'numeric'}, {'vector', 'nonempty'});
assert(all(regime == 1 | regime == 2), 'Regime entries must be 1 or 2.');
validateattributes(f, {'numeric'}, {'scalar', 'finite', '>', 1});
validateattributes(growth, {'numeric'}, {'positive', 'finite'});
validateattributes(weight, {'numeric'}, {'positive', 'finite'});

log2W = numel(regime) .* log2(f) .* (growth - 1) ...
        + sum(regime == 2) .* log2(weight);
W = 2 .^ log2W;
end
```
