## Supplementary Mathematical Model for "Obligate multicellularity circumvents population genetic barriers to collective-level adaptation": plot_ASRegime_outputs.m

```
function plot_ASRegime_outputs(regimes, names, f, growth, weight, meanSurvival)
% Output survival alone, ancestor-equivalent isoclines alone, and their overlay.
gx = log2(growth);
sy = log2(weight);
colors = [239 243 255; 198 219 239; 158 202 225; 107 174 214; 49 130 189; 8 81 156] / 255;
blues = interp1(linspace(0,1,size(colors,1)), colors, linspace(0,1,256));
stems = {'Figure5_survival', 'Figure5_fitness_isoclines', 'Figure5_combined'};
fid = fopen('Figure5_isoclines.csv', 'w');
assert(fid ~= -1, 'Cannot create isocline file.');
fprintf(fid, 'life_cycle,log2_growth_effect,log2_selection_effect,relative_cycle_fitness\n');
for r = 1:size(regimes,1)
    yy = linspace(0,3,601);
    base = 1 - sum(regimes(r,:) == 2) * yy / (numel(regimes(r,:)) * log2(f));
    keep = base >= 0.5 & base <= 2;
    xx = log2(base(keep));
    yy = yy(keep);
    W = cycle_fitness(regimes(r,:), f, 2.^xx, 2.^yy);
    for j = 1:numel(xx)
        fprintf(fid, '%s,%.17g,%.17g,%.17g\n', names{r}, xx(j), yy(j), W(j));
    end
end
fclose(fid);
for mode = 1:3
    fig = figure('Color', 'w', 'Position', [100 100 1000 850]);
    for r = 1:size(regimes,1)
        ax = subplot(2,2,r);
        if mode ~= 2
            imagesc(ax, gx, sy, meanSurvival(:,:,r)');
            axis(ax, 'xy');
            caxis(ax, [0 1]);
            colormap(ax, blues);
        end
        hold(ax, 'on');
        if mode ~= 1
            nM = sum(regimes(r,:) == 2);
            if nM == 0
                plot(ax, [0 0], [0 3], 'Color', [0.25 0.65 0.12], 'LineWidth', 2);
            else
                yy = linspace(0,3,601);
                base = 1 - nM * yy / (numel(regimes(r,:)) * log2(f));
                keep = base >= 0.5 & base <= 2;
                xx = log2(base(keep));
                plot(ax, xx, yy(keep), 'Color', [0.25 0.65 0.12], 'LineWidth', 2);
            end
        end
        xlim(ax, [-1.125 1.125]); ylim(ax, [-0.5 3.5]);
        set(ax, 'XTick', -1:0.25:1, 'YTick', 0:3, 'FontSize', 10, 'Box', 'off');
        title(ax, names{r});
        xlabel(ax, 'log_2 growth-rate multiplier');
        ylabel(ax, 'log_2 group-selection weight');
        if mode == 2
            text(ax, 0.25, 2.5, 'W > 1');
            text(ax, -0.95, 0.4, 'W < 1');
        end
    end
    if mode ~= 2
        cb = colorbar;
        ylabel(cb, 'Survival after eight rounds');
    end
    print(fig, [stems{mode}, '.png'], '-dpng', '-r300');
    print(fig, [stems{mode}, '.pdf'], '-dpdf', '-bestfit');
    close(fig);
end
end
```
