## Supplementary Mathematical Model for "Obligate multicellularity circumvents population genetic barriers to collective-level adaptation": run_ASRegime.m

```
% Generate survival estimates and relative-cycle fitness on the Figure 5 grid.
% ASregime performs 1000 stochastic runs per call.
% Requires MATLAB Statistics and Machine Learning Toolbox.

f = 32;
N = 1e6;
batches = 3;
seed = 20260917;
rng(seed, 'twister');
startTime = tic;

regimes = [2 2 2 2 2 2 2 2; 1 1 1 1 1 1 1 1; ...
           1 1 2 2 2 2 2 2; 1 1 1 1 2 2 2 2];
names = {'M100', 'U100', 'U25/M75', 'U50/M50'};
growth = 2 .^ (-1:0.25:1);
weight = [1 2 4 8];
counts = zeros(numel(growth), numel(weight), size(regimes, 1), batches);
W = zeros(numel(growth), numel(weight), size(regimes, 1));
log2W = W;

for r = 1:size(regimes, 1)
    for g = 1:numel(growth)
        for s = 1:numel(weight)
            [W(g,s,r), log2W(g,s,r)] = cycle_fitness(regimes(r,:), f, growth(g), weight(s));
            for b = 1:batches
                fprintf('%s: growth %d/9, selection %d/4, batch %d/3\n', names{r}, g, s, b);
                counts(g,s,r,b) = ASregime(regimes(r,:), f, N, growth(g), weight(s));
            end
        end
    end
end

meanSurvival = mean(counts, 4) / 1000;
fid = fopen('Figure5_results.csv', 'w');
assert(fid ~= -1, 'Cannot create results file.');
fprintf(fid, 'life_cycle,log2_growth_effect,log2_selection_effect,batch,survival_count_of_1000,relative_cycle_fitness,log2_relative_cycle_fitness\n');
for r = 1:size(regimes, 1)
    for g = 1:numel(growth)
        for s = 1:numel(weight)
            for b = 1:batches
                fprintf(fid, '%s,%.2f,%d,%d,%d,%.17g,%.17g\n', names{r}, log2(growth(g)), ...
                        log2(weight(s)), b, counts(g,s,r,b), W(g,s,r), log2W(g,s,r));
            end
        end
    end
end
fclose(fid);
environment = ver;
elapsedSeconds = toc(startTime);
save('Figure5_results.mat', 'regimes', 'names', 'growth', 'weight', 'f', 'N', ...
     'batches', 'seed', 'counts', 'meanSurvival', 'W', 'log2W', 'environment', 'elapsedSeconds');
plot_ASRegime_outputs(regimes, names, f, growth, weight, meanSurvival);
fprintf('Wrote survival, fitness and plot outputs in %.1f seconds.\n', elapsedSeconds);
```
