## Supplementary Mathematical Model for "Obligate multicellularity circumvents population genetic barriers to collective-level adaptation": plot_figure5.py

```
#!/usr/bin/env python3
"""Plot stochastic mutant survival and ancestor-equivalent fitness isoclines."""

import argparse
import csv
import hashlib
import json
from pathlib import Path

import matplotlib
matplotlib.use('Agg')
import matplotlib.pyplot as plt
from matplotlib.colors import LinearSegmentedColormap
import numpy as np

REGIMES = {
    'M100': [2]*8,
    'U100': [1]*8,
    'U25/M75': [1]*2+[2]*6,
    'U50/M50': [1]*4+[2]*4,
}


def cycle_fitness(regime, expansion, growth, weight):
    """Return relative fitness and its log2 value in the rare-mutant limit.

    Resident-determined growth time is ln(expansion). Relative abundance
    changes by expansion**(growth-1) per round and by weight per M transfer.
    This deterministic fitness calculation excludes discrete-group rounding,
    stochastic lineage loss and saturation at high mutant frequency.
    """
    if expansion <= 1 or not np.isfinite(expansion):
        raise ValueError('Population expansion must be finite and greater than 1.')
    if not regime or any(p not in (1, 2) for p in regime):
        raise ValueError('A regime must contain unicellular (1) or multicellular (2) rounds.')
    growth, weight = np.asarray(growth), np.asarray(weight)
    if not (np.all(np.isfinite(growth)) and np.all(growth > 0)
            and np.all(np.isfinite(weight)) and np.all(weight > 0)):
        raise ValueError('Growth multipliers and selection weights must be positive and finite.')
    log2_w = len(regime)*np.log2(expansion)*(growth-1) + regime.count(2)*np.log2(weight)
    return np.exp2(log2_w), log2_w


def read_survival(path, mode):
    counts = {}
    with path.open() as handle:
        rows = list(csv.DictReader(handle))
    for row in rows:
        key = (row['life_cycle'], float(row['log2_growth_effect']), float(row['log2_selection_effect']))
        if mode == 'means':
            if key in counts:
                raise ValueError('Duplicate parameter combination.')
            batches = {b: int(row[f'batch_{b}_count']) for b in (1,2,3)}
            np.testing.assert_allclose(np.mean(list(batches.values())), float(row['mean_survival_count']), rtol=0, atol=1e-10)
            counts[key] = batches
        else:
            batch = int(row['batch'])
            if batch in counts.setdefault(key, {}):
                raise ValueError('Duplicate batch.')
            counts[key][batch] = int(row['survival_count_of_1000'])
    expected = {(name, float(x), float(y)) for name in REGIMES
                for x in np.arange(-1,1.01,.25) for y in range(4)}
    if set(counts) != expected:
        raise ValueError('Input must contain the complete four-regime, 9-by-4 grid.')
    values = np.zeros((4,4,9))
    for i, name in enumerate(REGIMES):
        for y in range(4):
            for j, x in enumerate(np.arange(-1,1.01,.25)):
                batches = counts[name, x, y]
                if set(batches) != {1,2,3} or any(v < 0 or v > 1000 for v in batches.values()):
                    raise ValueError('Each combination needs three counts between 0 and 1000.')
                values[i,y,j] = np.mean(list(batches.values()))/1000
    return values


def isocline(regime, expansion):
    y = np.linspace(0, 3, 601)
    if regime.count(2) == 0:
        return np.zeros_like(y), y
    # Solve log2(W) = 0 for x = log2(growth-rate multiplier).
    growth = 1-regime.count(2)*y/(len(regime)*np.log2(expansion))
    keep = (growth >= .5) & (growth <= 2)
    return np.log2(growth[keep]), y[keep]


def draw(values, expansion, output, mode):
    plt.rcParams.update({'font.family':'Arial', 'font.size':10, 'svg.fonttype':'none',
                         'svg.hashsalt':'figure5-survival-fitness'})
    cmap = LinearSegmentedColormap.from_list('survival_blues',
        ['#eff3ff','#c6dbef','#9ecae1','#6baed6','#3182bd','#08519c'])
    fig, axes = plt.subplots(2,2,figsize=(7.15,6.8),sharex=True,sharey=True)
    fig.subplots_adjust(left=.10,right=.985,bottom=.235,top=.94,wspace=.07,hspace=.23)
    for i, ((name, regime), ax) in enumerate(zip(REGIMES.items(), axes.flat)):
        if mode != 'fitness_isoclines':
            mesh = ax.pcolormesh(np.arange(-1.125,1.126,.25), np.arange(-.5,4,1), values[i],
                                cmap=cmap,vmin=0,vmax=1,edgecolors='none',antialiased=False)
        else:
            xs, ys = np.linspace(-1,1,401), np.linspace(0,3,401)
            xx, yy = np.meshgrid(xs,ys)
            _, lw = cycle_fitness(regime, expansion, 2**xx, 2**yy)
            ax.contourf(xx,yy,lw,levels=[-1e6,0,1e6],colors=['white','#e7f3e2'])
            ax.text(.25,2.5,'Higher fitness',fontsize=9)
            ax.text(-.97,.4,'Lower fitness',fontsize=9)
        if mode != 'survival':
            xs, ys = isocline(regime, expansion)
            ax.plot(xs,ys,color='#50ad25',lw=1.7)
        ax.set_xlim(-1.125,1.125); ax.set_ylim(-.5,3.5)
        ax.set_title({'M100':'Multicellular (M100)','U100':'Unicellular (U100)'}.get(name,name),pad=8,fontsize=11)
        ax.text(-.04,1.055,chr(65+i),transform=ax.transAxes,fontsize=15,va='bottom')
        ax.set_xticks(np.arange(-1,1.01,.25))
        ax.set_xticklabels(['−1','−0.75','−0.5','−0.25','0','0.25','0.5','0.75','1'],fontsize=8)
        ax.set_yticks([0,1,2,3]); ax.tick_params(length=3,color='#c8c8c8',pad=4)
        for spine in ax.spines.values():
            spine.set_visible(False)
    fig.text(.015,.585,r'Group-selection weight ($\log_2$)',rotation=90,va='center',fontsize=11)
    fig.text(.54,.16,r'Growth-rate multiplier ($\log_2$)',ha='center',fontsize=11)
    if mode != 'fitness_isoclines':
        cax = fig.add_axes([.34,.092,.49,.025])
        cb = fig.colorbar(mesh,cax=cax,orientation='horizontal',ticks=np.linspace(0,1,6))
        cb.outline.set_visible(False); cb.ax.tick_params(length=2,labelsize=8)
        fig.text(.10,.105,'Survival after\neight rounds',va='center',fontsize=10)
    stem = 'Figure_5' if mode == 'combined' else f'Figure_5_{mode}'
    fig.savefig(output/f'{stem}.png',dpi=600)
    fig.savefig(output/f'{stem}.pdf',metadata={'CreationDate':None,'ModDate':None})
    fig.savefig(output/f'{stem}.svg',metadata={'Date':None})
    plt.close(fig)


def run(source, mode, expansion, output):
    values = read_survival(source, mode)
    output.mkdir(parents=True,exist_ok=True)
    for kind in ('survival','fitness_isoclines','combined'):
        draw(values, expansion, output, kind)
    with (output/'Figure_5_survival_and_fitness.csv').open('w',newline='') as handle:
        writer = csv.writer(handle)
        writer.writerow(['life_cycle','log2_growth_effect','log2_selection_effect','survival_fraction',
                         'relative_cycle_fitness','log2_relative_cycle_fitness'])
        for i,(name,regime) in enumerate(REGIMES.items()):
            for y in range(4):
                for j,x in enumerate(np.arange(-1,1.01,.25)):
                    w,lw = cycle_fitness(regime,expansion,2**x,2**y)
                    writer.writerow([name,x,y,values[i,y,j],float(w),float(lw)])
    max_error = 0
    with (output/'Figure_5_isocline_coordinates.csv').open('w',newline='') as handle:
        writer = csv.writer(handle)
        writer.writerow(['life_cycle','log2_growth_effect','log2_selection_effect','relative_cycle_fitness'])
        for name,regime in REGIMES.items():
            x,y = isocline(regime,expansion)
            w,lw = cycle_fitness(regime,expansion,2**x,2**y)
            max_error = max(max_error,float(np.max(np.abs(lw))))
            np.testing.assert_allclose(lw,0,rtol=0,atol=1e-12)
            writer.writerows(zip([name]*len(x),x,y,w))
    metadata = {
        'source':source.name,'source_sha256':hashlib.sha256(source.read_bytes()).hexdigest(),
        'survival_source_mode':mode,'population_expansion_per_round':expansion,
        'regimes':REGIMES,'survival_endpoint':'Mutant lineage present after eight rounds',
        'batches':3,'runs_per_batch':1000,'fitness_reference':'Ancestor without either mutation effect; W=1',
        'fitness_definition':'Deterministic relative abundance multiplier across a cycle in the rare-mutant limit',
        'log2_fitness_formula':'len(regime)*log2(expansion)*(growth_multiplier-1) + nM*log2(group_weight)',
        'fitness_assumptions':['Resident-determined growth interval ln(expansion)',
                               'Group weights multiply relative abundance in M rounds',
                               'No stochastic loss, group-count rounding or finite-frequency saturation in fitness calculation'],
        'green_isoclines':'Relative cycle fitness W = 1; higher W above, or to the right in U100',
        'isocline_max_abs_log2W':max_error,'native_MATLAB_execution':False,
        'numpy':np.__version__,'matplotlib':matplotlib.__version__,
    }
    (output/'Figure_5_output_record.json').write_text(json.dumps(metadata,indent=2)+'\n')
    print(f'Wrote three plot versions, survival/fitness values and isocline coordinates to {output}')


if __name__ == '__main__':
    parser = argparse.ArgumentParser(description=__doc__)
    inputs = parser.add_mutually_exclusive_group(required=True)
    inputs.add_argument('--mean-table',type=Path,help='Mean/count table from Supplementary Code 5')
    inputs.add_argument('--simulation-results',type=Path,help='Figure5_results.csv from the Python or MATLAB driver')
    parser.add_argument('--expansion',type=float,default=32)
    parser.add_argument('--output-dir',type=Path,required=True)
    args = parser.parse_args()
    run(args.mean_table or args.simulation_results, 'means' if args.mean_table else 'simulations',
        args.expansion,args.output_dir)
```
