## Supplementary Mathematical Model for "Obligate multicellularity circumvents population genetic barriers to collective-level adaptation": README.md

# Supplementary Code 2

This package simulates mutant-lineage survival through the four life cycles in Figure 5 and produces survival heatmaps, ancestor-equivalent fitness isoclines, and a combined figure. The package provides Python and MATLAB implementations.

## Run

Tested with Python 3.12.14 on macOS arm64. From the extracted package directory:

```sh
python -m pip install -r requirements.txt
python run_figure5.py --output-dir results
```

The default run uses seed 20260917 and all 144 combinations: four life cycles, nine growth-rate multipliers and four group-selection weights, each with three batches of 1,000 trials. Each parameter/batch combination has its own NumPy PCG64 stream derived from the master seed. A different seed can be supplied with `--seed`. No MATLAB installation or toolbox is required.

`results/Figure5_results.csv` contains the 432 batch counts and deterministic relative-fitness values. `Run_metadata.json` records exact parameters, random streams, runtime, software versions, source hashes and sampling diagnostics. The `figures` subdirectory contains survival-only, fitness-isocline-only and combined plots in PNG, PDF and SVG, plus numerical survival/fitness and isocline-coordinate tables.

## Plot Figure 5 data from the supplied counts

```sh
python plot_figure5.py --mean-table source/mean_survival_counts.csv --output-dir figure5_from_counts
```

`source/mean_survival_counts.csv` contains three survival counts per parameter combination in Figure 5. The same counts appear in the `figure5_survival_raw` worksheet of Supplementary_Data.xlsx. This code package runs without the workbook. Supplementary Code 5 analyzes the clustering shown in Supplementary Figure 4.

A new run gives another realization of the model. Its counts can differ from the Figure 5 counts because the simulation is stochastic. The same seed and pinned dependencies reproduce the supplied Python reference outputs. Plot appearance can depend on installed fonts; the supplied layout uses Arial.

## Model

Each trial starts with one mutant in a population of 1,000,000 cells. Growth expands the population 32-fold before transfer. The multicellular group size is 20 cells, as in `ASregime.m`. The schedules are eight M rounds, eight U rounds, two U followed by six M rounds, and four U followed by four M rounds. Growth-rate multipliers are calculated at full floating-point precision as 2 raised to log2 values from -1 to 1 in steps of 0.25; selection weights are 1, 2, 4 and 8.

`as_regime.py` uses a discrete growth-time grid, rounds mutant cell counts, converts mutant cells to groups by ceiling, and tracks survival after eight rounds. It searches the growth grid by binary search and advances replicates together. Floating-point threshold decisions and random draws can differ between Python and MATLAB.

Unweighted sampling uses hypergeometric draws or a finite-population normal approximation. Weighted sampling uses Poisson, binomial and high-frequency deterministic branches. The code reports an error for unsupported weighted-sampling conditions or an undefined growth-grid crossing.

Two edge-case rules apply: a negative rounded normal draw terminates a lineage as extinct, and a group draw at or above the available mutant-group count retains the pre-transfer mutant cell count, including any partial group. Both events are counted in run metadata.

`matlab_source/` contains the MATLAB model (`ASregime.m`), a driver for the Figure 5 grid (`run_ASRegime.m`) and the fitness and plotting functions.

## Fitness isoclines

Relative cycle fitness is calculated separately in the deterministic rare-mutant limit:

```text
W = f**(R*(g-1)) * w**nM
log2(W) = R*log2(f)*(g-1) + nM*log2(w)
```

Here R is the number of rounds, f the expansion factor, g the growth-rate multiplier, w the group-selection weight and nM the number of M rounds. The green curves mark W = 1. Fitness is higher above these curves; for U100 the boundary is vertical and fitness is higher to its right. The curved lines on log2 axes follow directly from this expression, as derived in Supplementary Note 1. Fitness excludes stochastic loss, discrete groups and finite-frequency saturation. It does not equal the probability of survival or fixation.
