## Supplementary Mathematical Model for "Obligate multicellularity circumvents population genetic barriers to collective-level adaptation": run_figure5.py

```
"""Run all 144 Figure 5 combinations, each in three batches of 1,000 trials."""

import argparse
from collections import Counter
import csv
import hashlib
import json
from pathlib import Path
import platform
import time

import numpy as np

from as_regime import simulate
from plot_figure5 import REGIMES, cycle_fitness, run as plot_results


def run(output, seed=20260917, plots=True):
    output.mkdir(parents=True, exist_ok=True)
    start = time.perf_counter()
    children = np.random.SeedSequence(seed).spawn(432)
    diagnostics = Counter()
    records = []
    for r, (name, regime) in enumerate(REGIMES.items()):
        for g, x in enumerate(np.arange(-1, 1.01, .25)):
            for y in range(4):
                growth, weight = float(2**x), float(2**y)
                fitness, log_fitness = cycle_fitness(regime, 32, growth, weight)
                for batch in range(1, 4):
                    stream = ((r*9+g)*4+y)*3+batch-1
                    result = simulate(regime, growth, weight,
                                      np.random.Generator(np.random.PCG64(children[stream])))
                    diagnostics.update(result.diagnostics)
                    records.append([name, float(x), y, batch, result.survival_count,
                                    float(fitness), float(log_fitness)])
            print(f'{name}: growth {g+1}/9 complete', flush=True)
    result_path = output/'Figure5_results.csv'
    with result_path.open('w', newline='') as handle:
        writer = csv.writer(handle)
        writer.writerow(['life_cycle', 'log2_growth_effect', 'log2_selection_effect', 'batch',
                         'survival_count_of_1000', 'relative_cycle_fitness', 'log2_relative_cycle_fitness'])
        writer.writerows(records)
    here = Path(__file__).resolve().parent
    metadata = {
        'seed': seed, 'rng': 'NumPy PCG64; SeedSequence child stream per parameter/batch',
        'stream_index': '((regime_index*9 + growth_index)*4 + log2_weight)*3 + batch_index',
        'population': 1000000, 'expansion': 32, 'group_size': 20, 'initial_mutants': 1,
        'runs_per_batch': 1000, 'batches': 3, 'parameter_combinations': 144,
        'total_trials': 432000, 'regimes': REGIMES,
        'growth_multipliers': [float(2**x) for x in np.arange(-1, 1.01, .25)],
        'selection_weights': [1, 2, 4, 8], 'python': platform.python_version(),
        'numpy': np.__version__, 'platform': platform.platform(),
        'elapsed_simulation_seconds': time.perf_counter()-start,
        'diagnostics': dict(diagnostics), 'native_MATLAB_execution': False,
        'source_sha256': {str(p.relative_to(here)): hashlib.sha256(p.read_bytes()).hexdigest()
                          for p in [here/'as_regime.py', here/'run_figure5.py',
                                    here/'matlab_source/ASregime.m']},
        'results_sha256': hashlib.sha256(result_path.read_bytes()).hexdigest(),
    }
    (output/'Run_metadata.json').write_text(json.dumps(metadata, indent=2)+'\n')
    if plots:
        plot_results(result_path, 'simulations', 32, output/'figures')
    print(f'Completed {len(records)*1000:,} trials in {metadata["elapsed_simulation_seconds"]:.1f} seconds.', flush=True)
    return metadata


if __name__ == '__main__':
    parser = argparse.ArgumentParser(description=__doc__)
    parser.add_argument('--output-dir', type=Path, required=True)
    parser.add_argument('--seed', type=int, default=20260917)
    parser.add_argument('--no-plots', action='store_true')
    args = parser.parse_args()
    run(args.output_dir, args.seed, not args.no_plots)
```
