## Supplementary Population Size for "Obligate multicellularity circumvents population genetic barriers to collective-level adaptation": build_figures.py

```
"""Draw population-size figures from the culture-level analysis outputs."""
import argparse
import csv
import json
from pathlib import Path
import numpy as np
import matplotlib
matplotlib.use('Agg')
import matplotlib.pyplot as plt
from matplotlib.lines import Line2D
from PIL import Image

TREATMENTS = ('U100', 'U50/M50', 'U25/M75', 'M100')
COLORS = ('#F7ABE8', '#55A0FB', '#C000C0', '#41F0AE')
DAY = np.arange(1, 9)

def significance_stars(p):
    return '****' if p < 0.0001 else '***' if p < 0.001 else '**' if p < 0.01 else '*' if p < 0.05 else 'ns'


def run(analysis_dir, assets_dir, output_dir):
    output_dir.mkdir(parents=True, exist_ok=True)
    result = json.loads((analysis_dir / 'results.json').read_text())
    rows = list(csv.DictReader((analysis_dir / 'population_sizes_long.csv').open()))
    y = np.full((8, 4, 3), np.nan)
    for row in rows:
        y[int(row['day'])-1, TREATMENTS.index(row['treatment']), int(row['replicate'])-1] = float(row['log10_population_size'])
    assert len(rows) == 96 and np.isfinite(y).all()
    p = [r['p_holm_three_primary_comparisons'] for r in result['primary_comparisons']]
    plt.rcParams.update({'font.family': 'DejaVu Sans', 'font.size': 13,
                         'axes.linewidth': 1.5, 'pdf.fonttype': 42, 'ps.fonttype': 42,
                         'svg.fonttype': 'none'})
    fig = plt.figure(figsize=(11, 5.6235), facecolor='white')
    ax = fig.add_axes([.075, .13, .56, .78])
    for g in (3, 2, 1, 0):
        ax.errorbar(DAY, y[:, g, :].mean(axis=1), yerr=y[:, g, :].std(axis=1, ddof=1),
                    color=COLORS[g], marker='o', markersize=7, linewidth=2.5,
                    elinewidth=1.2, capsize=4, label=TREATMENTS[g], zorder=3+g)
    baseline = [y[:,g,:].mean() for g in (0,3)]
    for value, color in zip(baseline, [COLORS[0],COLORS[3]]):
        ax.axhline(value, color=color, linewidth=2, linestyle=':', zorder=1)
    for x, color, pv in [(2, COLORS[2], p[2]), (4, COLORS[1], p[1])]:
        ax.annotate('', xy=(x,9.54), xytext=(x,9.69),
                    arrowprops={'arrowstyle':'-|>', 'color':color, 'lw':2.5})
        ax.text(x,9.78,'− β-estradiol',ha='center',fontsize=10)
        ax.text(x,9.96,significance_stars(pv),ha='center',fontsize=11)
    ax.plot([8.45,8.7,8.7,8.45], [baseline[0],baseline[0],baseline[1],baseline[1]],
            color='black',lw=1.2,clip_on=False)
    ax.text(8.86,np.mean(baseline),significance_stars(p[0]),
            fontsize=11,rotation=90,va='center',ha='center',clip_on=False)
    ax.set(xlim=(.6,8.45),ylim=(6.35,10.18),xticks=DAY,yticks=[7,8,9,10],xlabel='Day',
           ylabel='log₁₀ population size per culture')
    ax.spines[['top','right']].set_visible(False)
    ax.tick_params(length=5,width=1.5)
    handles=[Line2D([0],[0],color=c,marker='o',lw=2) for c in COLORS]
    fig.legend(handles,TREATMENTS,loc='upper left',bbox_to_anchor=(.09,1.01),
               ncol=4,frameon=False,fontsize=11,handlelength=1.3,columnspacing=1.1)
    fig.text(.015,.975,'A',fontsize=23,fontweight='bold',va='top')
    # The microscopy pixels and embedded scale bars are used without image adjustment.
    for name, label, title, bottom in [('microscopy_B.png','B','+ β-estradiol',.505),
                                       ('microscopy_C.png','C','− β-estradiol',.055)]:
        image = Image.open(assets_dir/name)
        width=.227; height=width*11/image.width*image.height/5.6235
        a=fig.add_axes([.765,bottom,width,height]);a.imshow(image,interpolation='none');a.set_axis_off()
        fig.text(.735,bottom+height+.012,label,fontsize=23,fontweight='bold',ha='right',va='bottom')
        fig.text(.8785,bottom+height+.016,title,fontsize=14,ha='center',va='bottom')
    save(fig,output_dir,'Figure_2')
    plt.close(fig)

    fig, axes=plt.subplots(1,2,figsize=(10,4.7),sharex=True,sharey=True)
    fig.subplots_adjust(left=.095,right=.98,bottom=.22,top=.89,wspace=.15)
    for j,(g,bp) in enumerate([(1,5),(2,3)]):
        ax=axes[j];pre=DAY<bp
        ax.axvspan(.75,bp-.5,color=COLORS[0],alpha=.14)
        ax.axvspan(bp-.5,8.25,color=COLORS[3],alpha=.12)
        ax.axvline(bp-.5,color='.55',linestyle=':',lw=1)
        for i,marker in enumerate(['o','s','^']):
            ax.plot(DAY,y[:,g,i],color=COLORS[g],alpha=.62,lw=.9,marker=marker,ms=5)
        for mask in [pre,~pre]:
            xx=DAY[mask]
            ax.plot([xx.min(),xx.max()],[y[mask,g,:].mean()]*2,color='black',lw=2.4)
            # Averaging culture-specific fits equals fitting the balanced daily means.
            beta=np.polyfit(xx,y[mask,g,:].mean(axis=1),1)
            plot_x=np.array([xx.min(), bp if mask[0] else xx.max()])
            ax.plot(plot_x,np.polyval(beta,plot_x),'--',color='black',lw=1.6)
        ax.set(title=TREATMENTS[g],xlabel='Day',xlim=(.75,8.25),ylim=(6.05,9.75),xticks=DAY,yticks=[6.5,7.5,8.5,9.5])
        ax.spines[['top','right']].set_visible(False)
        ax.text(-.13,1.1,'AB'[j],transform=ax.transAxes,fontsize=19,fontweight='bold',va='top')
    axes[0].set_ylabel('log₁₀ population size per culture')
    handles=[Line2D([0],[0],color='.5',marker=m,lw=.9,label=f'Culture {i+1}') for i,m in enumerate(['o','s','^'])]
    handles += [Line2D([0],[0],color='black',lw=2.4,label='Phase mean'),Line2D([0],[0],color='black',ls='--',lw=1.6,label='Segmented trend')]
    fig.legend(handles=handles,loc='lower center',bbox_to_anchor=(.53,.01),ncol=5,frameon=False,fontsize=10,columnspacing=1.4,handlelength=2.2)
    save(fig,output_dir,'Supplementary_Figure_3')
    plt.close(fig)
    means=[{'treatment':t,'day':int(d),'mean_log10':float(y[k,g,:].mean()),'sd_log10':float(y[k,g,:].std(ddof=1))}
           for g,t in enumerate(TREATMENTS) for k,d in enumerate(DAY)]
    with (output_dir/'plotted_means_and_sd.csv').open('w',newline='') as f:
        w=csv.DictWriter(f,fieldnames=means[0].keys());w.writeheader();w.writerows(means)


def save(fig,output_dir,name):
    fig.savefig(output_dir/f'{name}.png',dpi=400,facecolor='white')
    fig.savefig(output_dir/f'{name}.pdf',facecolor='white',metadata={'Title':name.replace('_',' '),'Author':'Peterson et al.'})
    fig.savefig(output_dir/f'{name}.svg',facecolor='white')


if __name__=='__main__':
    parser=argparse.ArgumentParser(description=__doc__)
    parser.add_argument('--analysis-dir',type=Path,required=True)
    parser.add_argument('--assets-dir',type=Path,default=Path(__file__).resolve().parent/'assets')
    parser.add_argument('--output-dir',type=Path,required=True)
    args=parser.parse_args();run(args.analysis_dir,args.assets_dir,args.output_dir)
```
