## Supplementary Population Size for "Obligate multicellularity circumvents population genetic barriers to collective-level adaptation": README.md

# Population-size analyses

This package reproduces the population-size comparisons in Figure 2 and Supplementary Figure 3 of Peterson et al., “Obligate multicellularity circumvents population genetic barriers to collective-level adaptation.”

The input is the `figure2_population_size` sheet, cells A2:M10, in `Supplementary_Data.xlsx`. It contains 96 observations from the same three cultures per treatment followed across eight days. Counts are units of selection per culture: single cells during unicellular growth and clusters during multicellular growth. No observations are excluded or rescaled.

## Run

Python 3.12 was used. Install the versions listed in `requirements.txt`, then run these commands from this directory. The example assumes the workbook is one directory above this folder; change the input path if needed.

```sh
python -m pip install -r requirements.txt
python reanalyze_population_sizes.py --workbook ../Supplementary_Data.xlsx --output-dir results
python build_figures.py --analysis-dir results --output-dir figures
```

The analysis writes culture summaries, fitted coefficients, all 96 observations with source-cell addresses, statistical results, and a diagnostic plot. The plotting script writes Figure 2 and Supplementary Figure 3 as PDF, PNG and SVG, plus the 32 plotted means and sample SDs. Figure 2 micrographs, including their scale bars, are in `assets/`.

## Comparisons

We use one eight-day mean log10 population size per culture for the U100 versus M100 Welch test. Facultative comparisons use each culture’s pre-minus-post phase means, equivalent to testing the coefficients from culture-specific piecewise-constant regressions. All tests are two-sided. The three primary p values are adjusted together using Holm’s method. Ratio confidence intervals are pointwise 95% t intervals, calculated on the log10 scale and back-transformed; they are not adjusted for multiple comparisons.

The segmented sensitivity analysis fits separate pre-switch and post-switch trends to each culture and tests their separation at the first multicellular measurement, day 5 for U50/M50 and day 3 for U25/M75. Its p values and intervals are unadjusted. The U25/M75 separation is uncertain because only two observations determine each culture’s pre-switch trend. This contrast differs from the phase-average comparison and does not measure an instantaneous change at estradiol removal.

The fixed schedule links phase to elapsed time. Culture-level t inference assumes approximately normal summaries or contrasts, which cannot be assessed reliably with three cultures.

## Reference output

`reference_results/` contains numerical outputs for the supplied workbook. Its SHA-256 is recorded in `results.json`. Key Holm-adjusted p values are 0.0007999653, 0.0333030783 and 0.0010822935 for U100/M100, U50/M50 and U25/M75, respectively. The segmented sensitivity p values are 0.0264728280 and 0.0928872866.
