## Supplementary Population Size for "Obligate multicellularity circumvents population genetic barriers to collective-level adaptation": reanalyze_population_sizes.py

```
"""Culture-level analyses of the eight-day population-size experiment.

Each culture is an independent replicate. Daily observations are retained within
cultures for estimation and plotting, but not counted as independent replicates.
"""

import argparse
import csv
import hashlib
import json
import platform
from pathlib import Path

import numpy as np
import openpyxl
import scipy
from scipy import stats


TREATMENTS = ("U100", "U50/M50", "U25/M75", "M100")
DAY = np.arange(1, 9, dtype=float)
BREAKPOINTS = {"U50/M50": 5, "U25/M75": 3}


def write_csv(path, rows):
    with path.open("w", newline="") as handle:
        writer = csv.DictWriter(handle, fieldnames=list(rows[0]))
        writer.writeheader()
        writer.writerows(rows)


def ols(x, y):
    coefficients, _, rank, _ = np.linalg.lstsq(x, y, rcond=None)
    assert rank == x.shape[1]
    residuals = y - x @ coefficients
    sse = float(residuals @ residuals)
    sst = float((y - y.mean()) @ (y - y.mean()))
    return coefficients, residuals, sse, float(1 - sse / sst)


def contrast_result(values):
    values = np.asarray(values, dtype=float)
    n = values.size
    mean = float(values.mean())
    se = float(values.std(ddof=1) / np.sqrt(n))
    t = mean / se
    df = n - 1
    p = float(2 * stats.t.sf(abs(t), df))
    interval = mean + np.array([-1, 1]) * stats.t.ppf(0.975, df) * se
    independent_check = stats.ttest_1samp(values, 0)
    np.testing.assert_allclose([t, p], [independent_check.statistic,
                                      independent_check.pvalue], rtol=1e-12)
    return {
        "independent_cultures": n,
        "estimate_log10": mean,
        "standard_error_log10": se,
        "t": float(t), "df": df, "p_two_sided": p,
        "ci95_log10_low": float(interval[0]),
        "ci95_log10_high": float(interval[1]),
        "ratio": float(10 ** mean),
        "ci95_ratio_low": float(10 ** interval[0]),
        "ci95_ratio_high": float(10 ** interval[1]),
    }


def run(workbook, output):
    output.mkdir(parents=True, exist_ok=True)
    input_hash = hashlib.sha256(workbook.read_bytes()).hexdigest()
    book = openpyxl.load_workbook(workbook, read_only=True, data_only=True)
    sheet = book["figure2_population_size"]
    assert sheet["A1"].value == "N = population size per culture"
    headers = [sheet.cell(2, c).value for c in range(2, 14)]
    assert headers == [f"{t}_{r}" for t in TREATMENTS for r in range(1, 4)]
    assert [sheet.cell(r, 1).value for r in range(3, 11)] == list(range(1, 9))
    counts = np.array([[sheet.cell(r, c).value for c in range(2, 14)]
                       for r in range(3, 11)], dtype=float).reshape(8, 4, 3)
    assert counts.size == 96 and np.isfinite(counts).all() and (counts > 0).all()
    y = np.log10(counts)
    long_rows = []
    for d, day in enumerate(DAY.astype(int)):
        for g, treatment in enumerate(TREATMENTS):
            for replicate in range(3):
                phase = ("unicellular" if treatment == "U100" or
                         (treatment in BREAKPOINTS and day < BREAKPOINTS[treatment])
                         else "multicellular")
                long_rows.append({"culture_id": f"{treatment}_{replicate + 1}",
                                  "treatment": treatment, "replicate": replicate + 1,
                                  "day": int(day), "phase": phase,
                                  "population_size_per_culture": float(counts[d, g, replicate]),
                                  "log10_population_size": float(y[d, g, replicate]),
                                  "source_cell": f"{openpyxl.utils.get_column_letter(2 + 3*g + replicate)}{day + 2}"})
    write_csv(output / "population_sizes_long.csv", long_rows)

    culture_rows = []
    primary = []
    segmented_results = []
    regressions = []

    # Independent obligate cultures: one summary per culture, not per day.
    u = y[:, 0, :].mean(axis=0)
    m = y[:, 3, :].mean(axis=0)
    test = stats.ttest_ind(u, m, equal_var=False)
    interval = test.confidence_interval()
    difference = float(u.mean() - m.mean())
    se = np.sqrt(u.var(ddof=1)/3 + m.var(ddof=1)/3)
    df_check = (u.var(ddof=1)/3 + m.var(ddof=1)/3)**2 / (
        (u.var(ddof=1)/3)**2/2 + (m.var(ddof=1)/3)**2/2)
    np.testing.assert_allclose([test.statistic, test.df], [difference/se, df_check])
    group_x = np.column_stack([np.ones(6), np.r_[np.ones(3), np.zeros(3)]])
    group_beta, _, _, group_r2 = ols(group_x, np.r_[u, m])
    np.testing.assert_allclose(group_beta[1], difference)
    primary.append({"comparison": "U100 vs M100", "estimand": "Eight-day culture mean log10 difference",
                    "independent_cultures": 6, "estimate_log10": difference,
                    "standard_error_log10": float(se), "t": float(test.statistic),
                    "df": float(test.df), "p_two_sided": float(test.pvalue),
                    "ci95_log10_low": float(interval.low), "ci95_log10_high": float(interval.high),
                    "ratio": float(10**difference), "ci95_ratio_low": float(10**interval.low),
                    "ci95_ratio_high": float(10**interval.high)})
    for treatment, values in [("U100", u), ("M100", m)]:
        for i, value in enumerate(values):
            culture_rows.append({"culture_id": f"{treatment}_{i+1}", "summary": "whole_cycle_mean_log10",
                                 "value": float(value), "back_transformed": float(10**value)})
    for treatment, breakpoint in BREAKPOINTS.items():
        g = TREATMENTS.index(treatment)
        pre = DAY < breakpoint
        post = (~pre).astype(float)
        centered_day = DAY - breakpoint
        x_const = np.column_stack([np.ones(8), post])
        x_segment = np.column_stack([np.ones(8), centered_day, post, centered_day*post])
        phase_contrasts = []
        line_contrasts = []
        segment_betas = []
        for i in range(3):
            observed = y[:, g, i]
            bc, rc, _, r2c = ols(x_const, observed)
            bs, rs, _, r2s = ols(x_segment, observed)
            pre_mean = float(observed[pre].mean()); post_mean = float(observed[~pre].mean())
            contrast = pre_mean - post_mean
            np.testing.assert_allclose(-bc[1], contrast, atol=1e-13)
            phase_contrasts.append(contrast); line_contrasts.append(float(-bs[2]))
            segment_betas.append(bs)
            for summary, value in [("pre_mean_log10", pre_mean), ("post_mean_log10", post_mean),
                                   ("phase_pre_minus_post_log10", contrast),
                                   ("first_post_day_pre_minus_post_fitted_lines", float(-bs[2]))]:
                culture_rows.append({"culture_id": f"{treatment}_{i+1}", "summary": summary,
                                     "value": value, "back_transformed": float(10**value)})
            regressions.append({"treatment": treatment, "replicate": i+1,
                                "first_post_day": breakpoint,
                                "constant_pre_intercept": float(bc[0]),
                                "constant_post_change": float(bc[1]),
                                "constant_r_squared": r2c,
                                "segmented_pre_intercept_at_first_post_day": float(bs[0]),
                                "segmented_pre_slope": float(bs[1]),
                                "segmented_post_change": float(bs[2]),
                                "segmented_slope_change": float(bs[3]),
                                "segmented_post_slope": float(bs[1] + bs[3]),
                                "segmented_r_squared": r2s})
            # A uniform x10 conversion must alter the intercept alone.
            for x, beta, r2 in [(x_const, bc, r2c), (x_segment, bs, r2s)]:
                shifted, _, _, shifted_r2 = ols(x, observed+1)
                np.testing.assert_allclose(shifted[1:], beta[1:], atol=1e-13)
                np.testing.assert_allclose(shifted[0], beta[0]+1, atol=1e-13)
                np.testing.assert_allclose(shifted_r2, r2, atol=1e-13)
        result = contrast_result(phase_contrasts)
        direct = stats.ttest_rel(y[pre, g, :].mean(axis=0), y[~pre, g, :].mean(axis=0))
        np.testing.assert_allclose([result["t"], result["p_two_sided"]],
                                  [direct.statistic, direct.pvalue])
        primary.append({"comparison": treatment, "estimand": "Culture-specific unicellular-minus-multicellular phase mean",
                        **result})
        segmented_results.append({"comparison": treatment, "estimand": "Pre-minus-post fitted line separation at first post day",
                                  **contrast_result(line_contrasts),
                                  "culture_contrasts": line_contrasts,
                                  "mean_coefficients": np.mean(segment_betas, axis=0).tolist()})
    # Apply Holm's adjustment across the three primary contrasts.
    order = np.argsort([r["p_two_sided"] for r in primary]); previous=0.0
    for position, index in enumerate(order):
        adjusted=min(1.0, max(previous, (3-position)*primary[index]["p_two_sided"]))
        primary[index]["p_holm_three_primary_comparisons"] = adjusted
        previous=adjusted
    write_csv(output / "primary_comparisons.csv", primary)
    write_csv(output / "culture_summaries.csv", culture_rows)
    write_csv(output / "culture_regression_coefficients.csv", regressions)
    assert hashlib.sha256(workbook.read_bytes()).hexdigest() == input_hash
    summary = {
        "source_workbook": workbook.name, "source_sha256": input_hash,
        "sheet": "figure2_population_size", "source_range": "A2:M10",
        "n_observations": 96, "n_cultures": 12, "cultures_per_treatment": 3,
        "excluded_observations": [], "log_base": 10,
        "environment": {"python": platform.python_version(), "numpy": np.__version__,
                        "scipy": scipy.__version__, "openpyxl": openpyxl.__version__},
        "primary_comparisons": primary, "trend_sensitive_segmented_regressions": segmented_results}
    (output / "results.json").write_text(json.dumps(summary, indent=2)+"\n")
    plot_results(y, output)
    print(json.dumps({"primary": primary, "segmented": segmented_results}, indent=2))


def plot_results(y, output):
    import matplotlib
    matplotlib.use("Agg")
    import matplotlib.pyplot as plt
    colors = ["#A83487", "#2476C7", "#AC00A9", "#008D69"]
    fig, axes = plt.subplots(2, 2, figsize=(10, 7), sharex=True, sharey=True,
                             layout="constrained")
    for g, (treatment, ax) in enumerate(zip(TREATMENTS, axes.flat)):
        for i, style in enumerate(["o-", "s-", "^-"]):
            ax.plot(DAY, y[:,g,i], style, color=colors[g], alpha=.55,
                    markersize=4, linewidth=1, label=f"Culture {i+1}")
        ax.plot(DAY, y[:,g,:].mean(axis=1), color="black", linewidth=2,
                label="Daily mean of log counts")
        if treatment in BREAKPOINTS:
            bp=BREAKPOINTS[treatment];pre=DAY<bp
            ax.axvspan(.7,bp-.5,color=colors[g],alpha=.06)
            ax.axvline(bp-.5,color="0.55",linestyle=":",linewidth=1)
            for phase in [pre,~pre]:
                x=DAY[phase]
                fit=np.polyfit(x,y[phase,g,:].mean(axis=1),1)
                ax.plot(x,np.polyval(fit,x),"--",color="black",linewidth=1.5,
                        label="Segmented trend" if np.all(phase==pre) else None)
        ax.set_title(treatment,fontsize=12)
        ax.spines[["top","right"]].set_visible(False)
        ax.set_xticks(range(1,9));ax.set_xlim(.7,8.3);ax.set_ylim(6.3,9.75)
        ax.set_xlabel("Day");ax.set_ylabel("log₁₀ population size per culture")
    axes[0,0].legend(frameon=False,fontsize=8,loc="lower left")
    axes[0,1].legend(frameon=False,fontsize=8,loc="lower left")
    fig.suptitle("Population trajectories across the same three cultures",fontsize=14)
    fig.savefig(output/'culture_trajectories.png',dpi=180)
    fig.savefig(output/'culture_trajectories.pdf')
    plt.close(fig)


if __name__ == "__main__":
    parser=argparse.ArgumentParser(description=__doc__)
    default_workbook=Path(__file__).resolve().parent.parent/'Supplementary_Data.xlsx'
    parser.add_argument('--workbook',type=Path,default=default_workbook)
    parser.add_argument('--output-dir',type=Path,default=Path(__file__).resolve().parent/'results')
    args=parser.parse_args()
    run(args.workbook,args.output_dir)
```
