## Supplementary figures and images for "Obligate multicellularity circumvents population genetic barriers to collective-level adaptation"

### microscopy_B.png

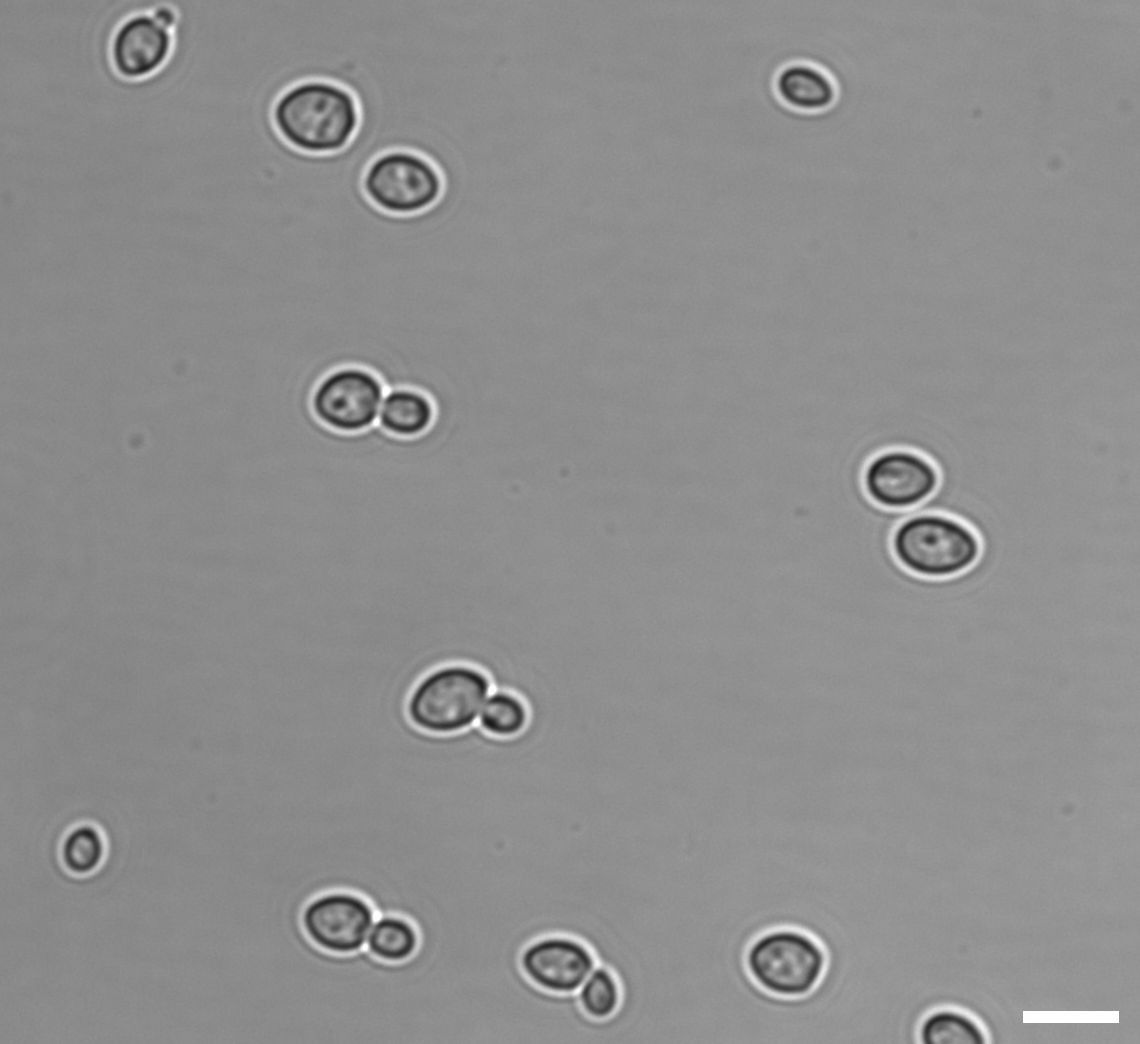

### microscopy_C.png

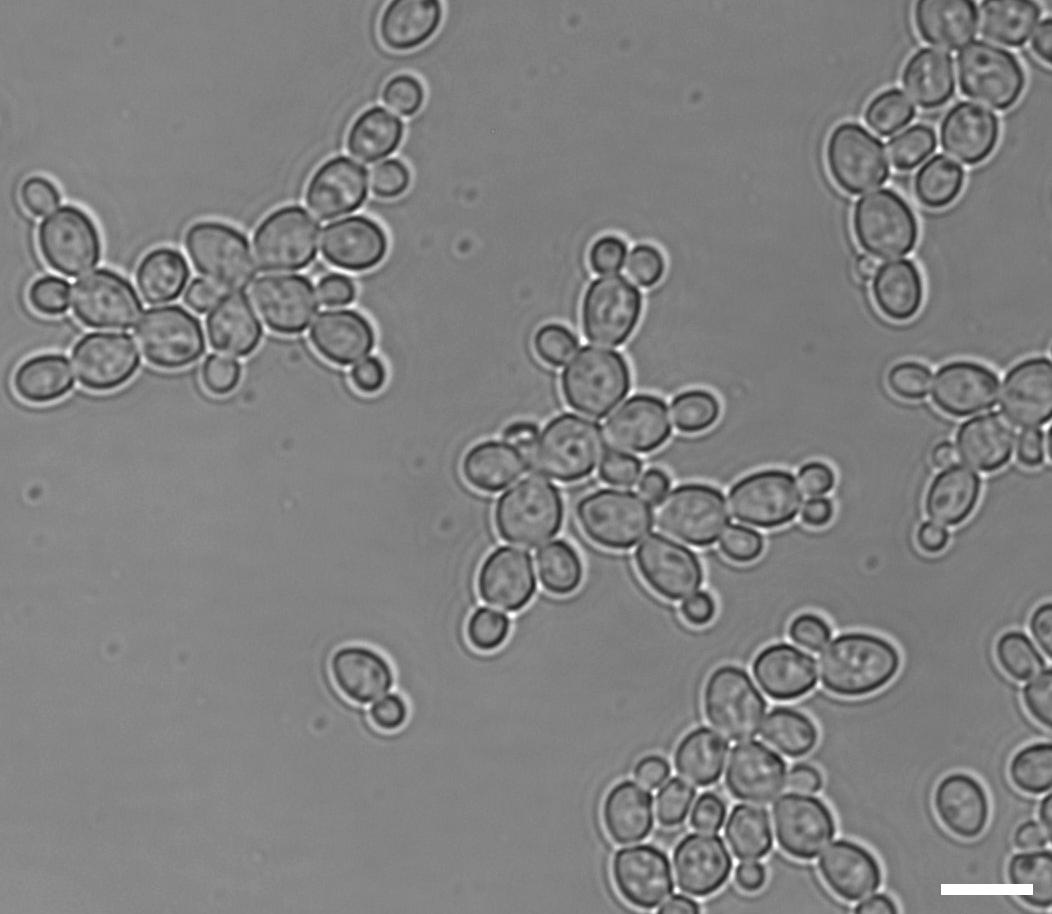

### panel_B.png

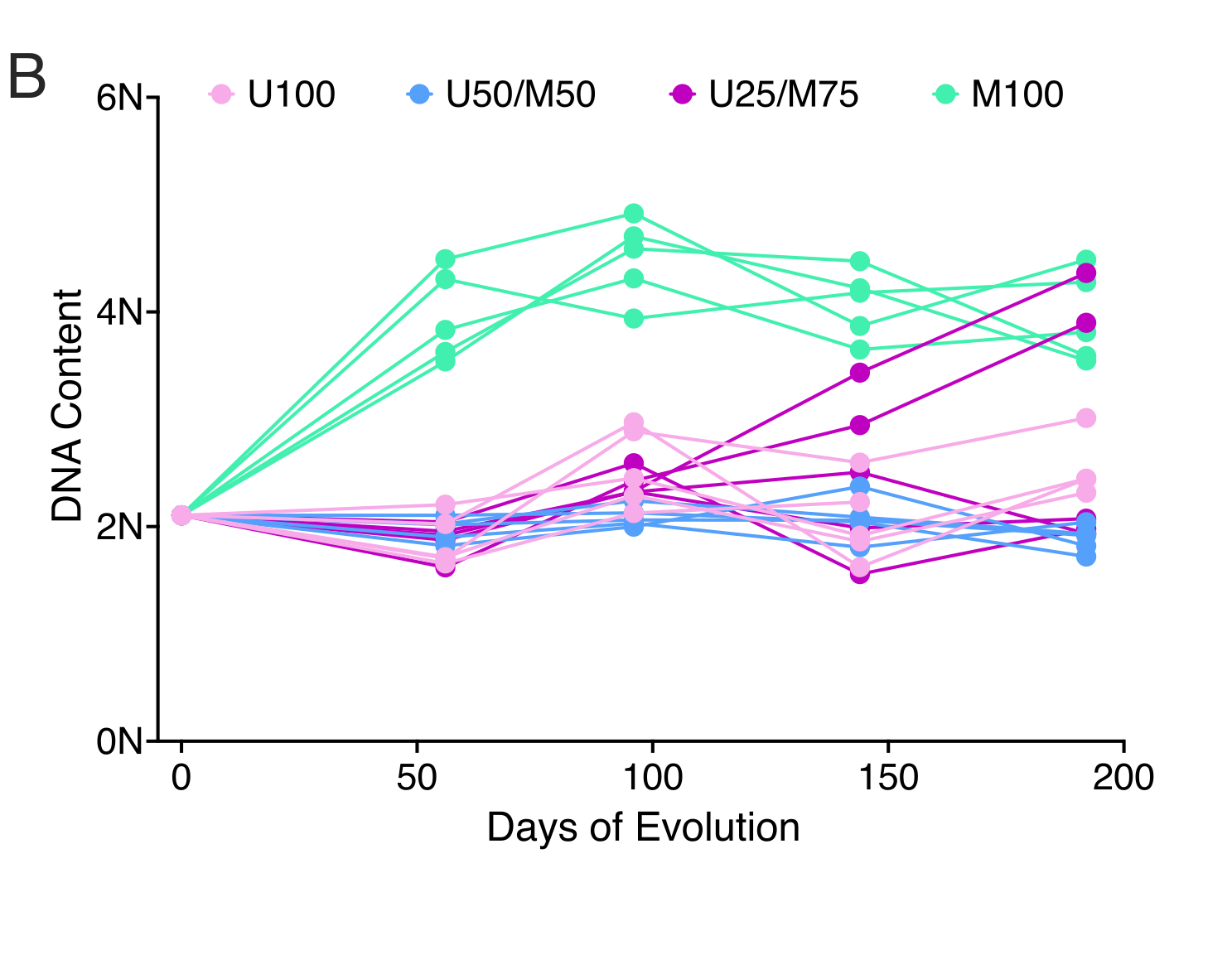
