## Supplementary Cluster Radius for "Obligate multicellularity circumvents population genetic barriers to collective-level adaptation": analyze_cluster_radius.py

```
"""Compare mean cluster radius among the three evolved multicellular treatments."""
import argparse
import csv
import hashlib
import itertools
import json
from pathlib import Path
import platform

import numpy as np
import openpyxl
import scipy
from scipy import stats

GROUPS = [('U50/M50', 7, 'fc50'), ('U25/M75', 12, 'f75'), ('M100', 17, 'multi')]


def write_csv(path, rows):
    with path.open('w', newline='') as handle:
        writer = csv.DictWriter(handle, fieldnames=list(rows[0]))
        writer.writeheader()
        writer.writerows(rows)


def run(workbook, output):
    output.mkdir(parents=True, exist_ok=True)
    original_hash = hashlib.sha256(workbook.read_bytes()).hexdigest()
    book = openpyxl.load_workbook(workbook, read_only=True, data_only=True)
    sheet = book['figure3_cluster_radius']
    assert sheet['A1'].value == 'ancestor'
    ancestor = float(sheet['B1'].value)
    groups, observations, summaries = [], [], []
    for treatment, first_row, prefix in GROUPS:
        values = []
        for r in range(first_row, first_row + 5):
            label = sheet.cell(r, 1).value.strip()
            assert label.startswith(prefix + ' ')
            value = float(sheet.cell(r, 2).value)
            values.append(value)
            observations.append({'treatment': treatment, 'population_label': label,
                                 'mean_radius_um': value, 'source_cell': f'B{r}'})
        x = np.array(values)
        assert np.isfinite(x).all() and (x > 0).all()
        groups.append(x)
        summaries.append({'treatment': treatment, 'n_populations': len(x),
                          'mean_radius_um': float(x.mean()),
                          'sd_population_means_um': float(x.std(ddof=1)),
                          'variance_population_means': float(x.var(ddof=1)),
                          'percent_above_ancestor_reference': float(100*(x.mean()/ancestor-1))})
    book.close()
    n = np.array([len(x) for x in groups])
    means = np.array([x.mean() for x in groups])
    variances = np.array([x.var(ddof=1) for x in groups])
    k = len(groups)
    grand_mean = np.average(means, weights=n)
    ss_between = float(np.sum(n*(means-grand_mean)**2))
    ss_within = float(np.sum((n-1)*variances))
    df_between, df_within = k-1, int(n.sum()-k)
    ms_between, ms_within = ss_between/df_between, ss_within/df_within
    anova = stats.f_oneway(*groups)
    np.testing.assert_allclose(anova.statistic, ms_between/ms_within, rtol=1e-12)
    np.testing.assert_allclose(anova.pvalue, stats.f.sf(anova.statistic, df_between, df_within), rtol=1e-12)
    tukey = stats.tukey_hsd(*groups)
    ci = tukey.confidence_interval(confidence_level=.95)
    pairwise = []
    for i, j in itertools.combinations(range(k), 2):
        # Report the later-listed treatment minus the earlier-listed treatment.
        se = np.sqrt(ms_within/2 * (1/n[i]+1/n[j]))
        q = abs(means[j]-means[i])/se
        expected_p = stats.studentized_range.sf(q, k, df_within)
        np.testing.assert_allclose(tukey.pvalue[j,i], expected_p, rtol=1e-12)
        pairwise.append({'comparison': f'{GROUPS[j][0]} minus {GROUPS[i][0]}',
                         'mean_difference_um': float(tukey.statistic[j,i]),
                         'p_tukey_family_adjusted': float(tukey.pvalue[j,i]),
                         'simultaneous_ci95_low_um': float(ci.low[j,i]),
                         'simultaneous_ci95_high_um': float(ci.high[j,i])})
    # Unequal-variance sensitivity analysis, motivated by the observed variance spread.
    welch = stats.f_oneway(*groups, equal_var=False)
    weights = n/variances
    weighted_mean = np.average(means, weights=weights)
    correction = np.sum((1-weights/weights.sum())**2/(n-1))
    welch_df = (k*k-1)/(3*correction)
    welch_f = np.sum(weights*(means-weighted_mean)**2)/(k-1)/(1+2*(k-2)*correction/(k*k-1))
    np.testing.assert_allclose(welch.statistic, welch_f, rtol=1e-12)
    np.testing.assert_allclose(welch.pvalue, stats.f.sf(welch_f, k-1, welch_df), rtol=1e-12)
    gh = stats.tukey_hsd(*groups, equal_var=False)
    gh_ci = gh.confidence_interval(confidence_level=.95)
    gh_pairs = []
    for i,j in itertools.combinations(range(k),2):
        a,b = variances[i]/n[i], variances[j]/n[j]
        df = (a+b)**2/(a*a/(n[i]-1)+b*b/(n[j]-1))
        expected_p = stats.studentized_range.sf(abs(means[j]-means[i])/np.sqrt((a+b)/2), k, df)
        np.testing.assert_allclose(gh.pvalue[j,i], expected_p, rtol=1e-12)
        gh_pairs.append({'comparison': f'{GROUPS[j][0]} minus {GROUPS[i][0]}',
                         'mean_difference_um': float(gh.statistic[j,i]),
                         'p_games_howell_family_adjusted': float(gh.pvalue[j,i]),
                         'ci95_low_um': float(gh_ci.low[j,i]),
                         'ci95_high_um': float(gh_ci.high[j,i])})
    assert hashlib.sha256(workbook.read_bytes()).hexdigest() == original_hash
    result = {'source_workbook': workbook.name, 'source_sha256': original_hash,
              'sheet': 'figure3_cluster_radius', 'evolved_data_range': 'A7:B21',
              'ancestor_reference': {'source_cell': 'B1', 'radius_um': ancestor,
                                     'used_for_inference': False,
                                     'note': 'Descriptive ancestor reference; excluded from all inferential tests.'},
              'group_summaries': summaries,
              'ordinary_one_way_anova': {'F': float(anova.statistic), 'df_numerator': df_between,
                                        'df_denominator': df_within, 'p': float(anova.pvalue),
                                        'ss_between': ss_between, 'ss_within': ss_within},
              'tukey_hsd': pairwise,
              'unequal_variance_sensitivity': {'sample_variance_ratio_max_to_min': float(variances.max()/variances.min()),
                  'welch_anova': {'F': float(welch.statistic), 'df_numerator': k-1,
                                  'df_denominator': float(welch_df), 'p': float(welch.pvalue)},
                  'games_howell': gh_pairs},
              'environment': {'python': platform.python_version(), 'numpy': np.__version__,
                              'scipy': scipy.__version__, 'openpyxl': openpyxl.__version__},
              }
    write_csv(output/'population_means.csv', observations)
    write_csv(output/'group_summaries.csv', summaries)
    write_csv(output/'tukey_comparisons.csv', pairwise)
    write_csv(output/'games_howell_sensitivity.csv', gh_pairs)
    (output/'results.json').write_text(json.dumps(result, indent=2)+'\n')
    print(json.dumps(result,indent=2))


if __name__ == '__main__':
    parser = argparse.ArgumentParser(description=__doc__)
    parser.add_argument('--workbook', type=Path, default=Path(__file__).resolve().parent.parent/'Supplementary_Data.xlsx')
    parser.add_argument('--output-dir', type=Path, default=Path(__file__).resolve().parent/'results')
    args = parser.parse_args()
    run(args.workbook, args.output_dir)
```
