## Supplementary Cluster Radius for "Obligate multicellularity circumvents population genetic barriers to collective-level adaptation": build_figure3.py

```
"""Draw Figure 3A and combine it with the Figure 3B artwork."""
import argparse
import csv
import hashlib
import json
from pathlib import Path
import xml.etree.ElementTree as ET

import matplotlib
matplotlib.use('Agg')
import matplotlib.pyplot as plt
import numpy as np
import openpyxl
from PIL import Image


def draw_panel(ax, values, adjusted_p):
    colors = ['black', '#F7ABE8', 'black', '#55A0FB', '#C000C0', '#41F0AE']
    labels = ['Anc_Uni', 'U100', 'Anc_Multi', 'U50/M50', 'U25/M75', 'M100']
    for x, (y, color) in enumerate(zip(values, colors)):
        offsets = np.zeros(len(y))
        placed = []
        for i in np.argsort(y):
            for candidate in [0, -.23, .23, -.115, .115]:
                if all(((candidate-offsets[j])/.21)**2 + ((y[i]-y[j])/.78)**2 >= 1 for j in placed):
                    offsets[i] = candidate
                    break
            else:
                raise ValueError('Point spacing requires a wider category')
            placed.append(i)
        ax.scatter(x+offsets, y, s=42, marker='s' if x < 2 else 'o',
                   color=color, edgecolors='none', zorder=3)
        ax.plot([x-.29, x+.29], [np.mean(y)]*2, color=color, lw=1.3)
    ax.plot([3,3,5,5], [24.0,24.7,24.7,24.0], color='black', lw=1.2)
    ax.text(4,25.15, ('*' if adjusted_p < 0.05 else 'ns'), ha='center', fontsize=11)
    ax.set(xlim=(-.65,5.65), ylim=(0,27), yticks=np.arange(0,26,5), ylabel='Radius (µm)')
    ax.set_xticks(range(6), labels, rotation=48, ha='right', rotation_mode='anchor')
    ax.tick_params(axis='x',pad=6)
    ax.spines[['top','right']].set_visible(False)
    ax.tick_params(length=4,width=1.3)


def run(workbook, analysis_dir, panel_b, output):
    output.mkdir(parents=True, exist_ok=True)
    result=json.loads((analysis_dir/'results.json').read_text())
    assert hashlib.sha256(workbook.read_bytes()).hexdigest()==result['source_sha256']
    book=openpyxl.load_workbook(workbook,read_only=True,data_only=True)
    s=book['figure3_cluster_radius']
    values=[np.array([s['B2'].value]), np.array([s.cell(r,2).value for r in range(3,7)]),
            np.array([s['B1'].value])]
    values += [np.array([s.cell(r,2).value for r in range(start,start+5)]) for start in [7,12,17]]
    for y, summary in zip(values[3:],result['group_summaries']):
        np.testing.assert_allclose(np.mean(y),summary['mean_radius_um'],rtol=1e-14)
    p=next(row['p_tukey_family_adjusted'] for row in result['tukey_hsd'] if row['comparison']=='M100 minus U50/M50')
    plt.rcParams.update({'font.family':'DejaVu Sans','font.size':13,'axes.linewidth':1.3,
                         'pdf.fonttype':42,'svg.fonttype':'none'})
    width,height=2517,1159
    dpi=width/11
    fig=plt.figure(figsize=(11,height/dpi),facecolor='white')
    ax=fig.add_axes([.096,.228,.297,.674])
    draw_panel(ax,values,p)
    fig.text(.005,.949,'A',fontsize=22,va='top')
    image_b=Image.open(panel_b).convert('RGB')
    assert image_b.size==(1487,1159)
    b=fig.add_axes([1030/width,0,1487/width,1])
    b.imshow(image_b,interpolation='none',aspect='auto');b.set_axis_off()
    fig.savefig(output/'Figure_3.pdf',facecolor='white',metadata={'Title':'Figure 3','Author':'Peterson et al.'})
    fig.savefig(output/'Figure_3.svg',facecolor='white')
    fig.savefig(output/'Figure_3.png',dpi=dpi,facecolor='white')
    plt.close(fig)
    vector_b = panel_b.with_suffix('.svg')
    if vector_b.exists():
        # Preserve the supplied panel's vector paths in the editable SVG export.
        namespace = 'http://www.w3.org/2000/svg'
        ET.register_namespace('', namespace)
        tree = ET.parse(output/'Figure_3.svg')
        panel = tree.find(f'.//{{{namespace}}}g[@id="axes_2"]')
        images = list(panel.iter(f'{{{namespace}}}image'))
        assert len(images) == 1
        raster = images[0]
        parent = next(e for e in panel.iter() if raster in list(e))
        position = list(parent).index(raster)
        vector = ET.parse(vector_b).getroot()
        vector.set('width', raster.get('width'))
        vector.set('height', raster.get('height'))
        group = ET.Element(f'{{{namespace}}}g', {'transform': raster.get('transform')})
        group.append(vector)
        parent.remove(raster)
        parent.insert(position, group)
        tree.write(output/'Figure_3.svg', encoding='utf-8', xml_declaration=True)
    # Embed the panel B artwork in the PNG composite.
    composite=Image.open(output/'Figure_3.png').convert('RGB')
    assert composite.size==(width,height),composite.size
    composite.paste(image_b,(1030,0));composite.save(output/'Figure_3.png',dpi=(dpi,dpi))
    assert np.array_equal(np.asarray(composite)[:,1030:,:],np.asarray(image_b))
    fig=plt.figure(figsize=(5,5.3),facecolor='white')
    ax=fig.add_axes([.16,.235,.82,.69]);draw_panel(ax,values,p)
    fig.savefig(output/'Figure_3A.pdf',facecolor='white',metadata={'Title':'Figure 3A','Author':'Peterson et al.'})
    fig.savefig(output/'Figure_3A.png',dpi=400,facecolor='white')
    fig.savefig(output/'Figure_3A.svg',facecolor='white');plt.close(fig)
    rows=[]
    for indices in [[2],list(range(3,7)),[1],list(range(7,12)),list(range(12,17)),list(range(17,22))]:
        for r in indices:rows.append({'label':str(s.cell(r,1).value).strip(),'mean_radius_um':s.cell(r,2).value,'source_cell':f'B{r}'})
    with (output/'plotted_values.csv').open('w',newline='') as f:
        writer=csv.DictWriter(f,fieldnames=rows[0].keys());writer.writeheader();writer.writerows(rows)
    book.close()


if __name__=='__main__':
    p=argparse.ArgumentParser(description=__doc__)
    p.add_argument('--workbook',type=Path,required=True)
    p.add_argument('--analysis-dir',type=Path,required=True)
    p.add_argument('--panel-b',type=Path,default=Path(__file__).resolve().parent/'assets/panel_B.png')
    p.add_argument('--output-dir',type=Path,required=True)
    args=p.parse_args();run(args.workbook,args.analysis_dir,args.panel_b,args.output_dir)
```
