## Supplementary Cluster Radius for "Obligate multicellularity circumvents population genetic barriers to collective-level adaptation": README.md

# Cluster-radius analyses

This package reproduces the Figure 3A analysis in Peterson et al., “Obligate multicellularity circumvents population genetic barriers to collective-level adaptation.”

The input is the `figure3_cluster_radius` sheet in `Supplementary_Data.xlsx`. Cells A7:B21 contain five population mean radii per evolved multicellular treatment: U50/M50, U25/M75 and M100. The sheet contains 15 population mean radii (µm). The population, rather than each measured cluster, is the unit of replication.

## Run

Python 3.12 was used. From this directory, run:

```sh
python -m pip install -r requirements.txt
python analyze_cluster_radius.py --workbook ../Supplementary_Data.xlsx --output-dir results
python build_figure3.py --workbook ../Supplementary_Data.xlsx --analysis-dir results --output-dir figures
```

Change the workbook path if the input is elsewhere. The analysis writes population means with source-cell addresses, group summaries, pairwise comparisons and `results.json`. The plotting script writes the combined Figure 3 and a separate Figure 3A as PNG, PDF and SVG, together with the plotted values. It reads the p label from the computed results.

## Analysis and interpretation

The primary analysis is a one-way ANOVA across the three evolved multicellular treatments, followed by Tukey’s HSD for all three pairs. Confidence intervals for these differences are simultaneous 95% intervals; Tukey p values account for the three pairwise comparisons. Welch’s ANOVA and Games–Howell comparisons provide a sensitivity analysis allowing unequal variances. Both approaches assume independent population means and rely on distributional approximations.

Ancestors and the four evolved U100 population means are displayed in panel A for context and excluded from inference. The multicellular ancestor value in B1 is used only as a descriptive reference for percentage differences. Horizontal offsets separate overlapping points and do not encode another measured variable. Horizontal bars show means.

The ordinary ANOVA gives F(2,12) = 4.07844, p = 0.0445192. Tukey-adjusted p values are 0.0372824 for M100 versus U50/M50, 0.236398 for M100 versus U25/M75 and 0.526612 for U25/M75 versus U50/M50. Welch/Games–Howell gives the same qualitative pattern. These analyses do not test an evolved-versus-ancestor difference or establish equivalence for the nonsignificant pairs.

## Figure preparation

The script draws panel A from the workbook and combines it with the panel B artwork in `assets/`. It writes the combined Figure 3 as PNG, PDF and SVG.

The data for panel B are in `figure3_ploidytimescales`. U100 population 3 has no day-192 value because its cells had begun to clump; its earlier measurements are included.

`reference_results/` contains the numerical outputs for the supplied workbook. The source hash and software versions are recorded in `results.json`.  
