## Supplementary Survival Clustering for "Obligate multicellularity circumvents population genetic barriers to collective-level adaptation": cluster_survival.py

```
#!/usr/bin/env python3
"""Reproduce Supplementary Figure 4 from mutant-survival counts."""

import argparse
import csv
import hashlib
import json
from pathlib import Path
import platform

import matplotlib
matplotlib.use("Agg")
import matplotlib.pyplot as plt
import numpy as np
import openpyxl
import scipy
from scipy.cluster.hierarchy import dendrogram, linkage
from scipy.spatial.distance import pdist, squareform


REGIMES = {
    "M100": "2,2,2,2,2,2,2,2",
    "U25/M75": "1,1,2,2,2,2,2,2",
    "U100": "1,1,1,1,1,1,1,1",
    "U50/M50": "1,1,1,1,2,2,2,2",
}
RAW_SHEET = "figure5_survival_raw"
SUMMARY_SHEET = "figure5_survival_summary"


def read_counts(path):
    """Read the four plotted regimes, retaining the source address of each count."""
    workbook = openpyxl.load_workbook(path, data_only=True, read_only=True)
    sheet = workbook[RAW_SHEET]
    expected_header = [
        "Regime (1=U, 2=M)", "Plotted treatment", "Batch", "Growth index",
        "Selection index", "log2 growth effect", "log2 selection effect",
        "Survivors after 8 rounds", "Runs per batch", "Survival fraction",
    ]
    if list(next(sheet.iter_rows(min_row=1, max_row=1, values_only=True))) != expected_header:
        raise ValueError("The survival-count worksheet layout has changed.")
    names = list(REGIMES)
    counts = np.full((4, 3, 9, 4), np.nan)
    cells = {}
    for row_number, row in enumerate(sheet.iter_rows(min_row=2, values_only=True), 2):
        if row[0] not in REGIMES.values():
            continue
        regime, name, batch, growth, selection, log_growth, log_selection, count, runs = row[:9]
        if name not in REGIMES or REGIMES[name] != regime:
            raise ValueError(f"Regime/name mismatch at row {row_number}.")
        if batch not in (1, 2, 3) or growth not in range(1, 10) or selection not in range(1, 5):
            raise ValueError(f"Unexpected matrix index at row {row_number}.")
        if log_growth != -1 + (growth - 1) / 4 or log_selection != selection - 1:
            raise ValueError(f"Parameter mapping mismatch at row {row_number}.")
        if runs != 1000 or not isinstance(count, (int, float)) or not np.isfinite(count):
            raise ValueError(f"Invalid survival count or run count at row {row_number}.")
        if not 0 <= count <= runs or count != int(count):
            raise ValueError(f"Survival count must be an integer from 0 to 1000, row {row_number}.")
        index = (names.index(name), batch - 1, growth - 1, selection - 1)
        if np.isfinite(counts[index]):
            raise ValueError(f"Duplicate regime/batch/parameter combination at row {row_number}.")
        counts[index] = count
        cells[index] = f"H{row_number}"
    if not np.isfinite(counts).all():
        raise ValueError("The four plotted regimes require 432 counts: 4 × 3 × 9 × 4.")

    # Check the workbook summaries against the raw counts.
    means = counts.mean(axis=1)
    seen = set()
    for row in workbook[SUMMARY_SHEET].iter_rows(min_row=2, values_only=True):
        if row[0] not in REGIMES:
            continue
        name, regime, lg, ls = row[:4]
        growth = int(round((lg + 1) * 4))
        selection = int(ls)
        key = (names.index(name), growth, selection)
        if key in seen or REGIMES[name] != regime:
            raise ValueError("Duplicate or mislabeled summary combination.")
        if not (0 <= growth < 9 and 0 <= selection < 4 and lg == -1 + growth / 4 and ls == selection):
            raise ValueError("Unexpected summary parameter combination.")
        np.testing.assert_allclose(row[4:7], counts[key[0], :, growth, selection], rtol=0, atol=0)
        np.testing.assert_allclose([row[7], row[8]], [means[key], means[key] / 1000], rtol=0, atol=1e-10)
        if row[9] != 1000:
            raise ValueError("Unexpected summary denominator.")
        seen.add(key)
    if len(seen) != 144:
        raise ValueError("The workbook summary must contain all 144 plotted combinations.")
    workbook.close()
    return names, counts, cells


def direct_ward(features):
    """Check Ward merges using the increase in within-cluster sum of squares."""
    clusters = {i: [i] for i in range(len(features))}
    result = []
    for step in range(len(features) - 1):
        choices = []
        ids = sorted(clusters)
        for j, left in enumerate(ids):
            for right in ids[j + 1:]:
                a, b = clusters[left], clusters[right]
                delta = len(a) * len(b) / (len(a) + len(b)) * np.sum(
                    (features[a].mean(axis=0) - features[b].mean(axis=0)) ** 2
                )
                choices.append((float(delta), left, right))
        delta, left, right = min(choices)
        members = clusters.pop(left) + clusters.pop(right)
        result.append([left, right, np.sqrt(2 * delta), len(members)])
        clusters[len(features) + step] = members
    return np.array(result)


def write_csv(path, header, rows):
    with path.open("w", newline="") as handle:
        writer = csv.writer(handle)
        writer.writerow(header)
        writer.writerows(rows)


def analyse(workbook_path, output):
    source_hash = hashlib.sha256(workbook_path.read_bytes()).hexdigest()
    names, counts, cells = read_counts(workbook_path)
    features = counts.mean(axis=1).reshape(4, 36)
    # Rows are life cycles, columns are matched parameter combinations. No standardization.
    tree = linkage(features, method="ward", metric="euclidean")
    np.testing.assert_allclose(tree, direct_ward(features), rtol=1e-12, atol=1e-10)
    scaled_tree = linkage(features / 1000, method="ward", metric="euclidean")
    np.testing.assert_array_equal(tree[:, [0, 1, 3]], scaled_tree[:, [0, 1, 3]])
    np.testing.assert_allclose(tree[:, 2] / 1000, scaled_tree[:, 2], rtol=1e-12)
    distances = squareform(pdist(features, metric="euclidean"))
    output.mkdir(parents=True, exist_ok=True)
    rows = []
    for i, name in enumerate(names):
        for growth in range(9):
            for selection in range(4):
                rows.append([name, -1 + growth / 4, selection,
                             *counts[i, :, growth, selection].astype(int).tolist(),
                             float(counts[i, :, growth, selection].mean()),
                             *[cells[i, batch, growth, selection] for batch in range(3)]])
    write_csv(output / "mean_survival_counts.csv", [
        "life_cycle", "log2_growth_effect", "log2_selection_effect",
        "batch_1_count", "batch_2_count", "batch_3_count", "mean_survival_count",
        "batch_1_source_cell", "batch_2_source_cell", "batch_3_source_cell"], rows)
    write_csv(output / "euclidean_distances.csv", ["life_cycle", *names],
              [[name, *row.tolist()] for name, row in zip(names, distances)])
    members = {i: [name] for i, name in enumerate(names)}
    merges = []
    for step, (left, right, height, number) in enumerate(tree):
        left, right = int(left), int(right)
        merges.append({"step": step + 1, "left": members[left], "right": members[right],
                       "height_count_scale": float(height), "n_life_cycles": int(number)})
        members[4 + step] = members[left] + members[right]
    write_csv(output / "ward_linkage.csv", ["step", "left_members", "right_members", "height_count_scale", "n_life_cycles"],
              [[m["step"], ";".join(m["left"]), ";".join(m["right"]), m["height_count_scale"], m["n_life_cycles"]] for m in merges])

    with plt.rc_context({"font.family": "DejaVu Sans", "font.size": 11,
                         "axes.linewidth": 1, "svg.hashsalt": "survival-clustering"}):
        fig, ax = plt.subplots(figsize=(7, 5))
        dendrogram(tree, labels=names, ax=ax, link_color_func=lambda _: "black", leaf_font_size=11)
        ax.set_ylabel("Ward linkage distance (survival-count scale)")
        ax.set_xlabel("Life cycle")
        ax.set_ylim(0, 2050)
        ax.spines[["top", "right"]].set_visible(False)
        fig.tight_layout()
        fig.savefig(output / "survival_clustering.png", dpi=300)
        fig.savefig(output / "survival_clustering.pdf", metadata={"CreationDate": None, "ModDate": None})
        fig.savefig(output / "survival_clustering.svg", metadata={"Date": None})
        plt.close(fig)
    if hashlib.sha256(workbook_path.read_bytes()).hexdigest() != source_hash:
        raise RuntimeError("Source workbook changed during analysis.")
    result = {
        "source_workbook": workbook_path.name, "source_sha256": source_hash,
        "raw_sheet": RAW_SHEET, "summary_sheet": SUMMARY_SHEET,
        "life_cycle_order": names, "features_per_life_cycle": 36,
        "batches_per_combination": 3, "runs_per_batch": 1000,
        "endpoint": "Mutant-lineage survival after eight rounds of growth and transfer",
        "feature_scale": "Mean survival counts out of 1000", "standardization": "none",
        "metric": "euclidean", "linkage": "ward", "merges": merges,
        "probability_scale_heights": scaled_tree[:, 2].tolist(),
        "environment": {"python": platform.python_version(), "numpy": np.__version__,
                        "scipy": scipy.__version__, "openpyxl": openpyxl.__version__,
                        "matplotlib": matplotlib.__version__},
    }
    (output / "results.json").write_text(json.dumps(result, indent=2) + "\n")
    return result


if __name__ == "__main__":
    parser = argparse.ArgumentParser(description=__doc__)
    parser.add_argument("--workbook", type=Path, required=True)
    parser.add_argument("--output-dir", type=Path, default=Path("results"))
    args = parser.parse_args()
    result = analyse(args.workbook, args.output_dir)
    for merge in result["merges"]:
        print(f"{merge['left']} + {merge['right']}: {merge['height_count_scale']:.6f}")
```
