## Supplementary Survival Clustering for "Obligate multicellularity circumvents population genetic barriers to collective-level adaptation": README.md

# Mutant-survival clustering

This package reproduces the hierarchical clustering in Supplementary Figure 4 of Peterson et al., “Obligate multicellularity circumvents population genetic barriers to collective-level adaptation.”

## Run

Python 3.12 was used. Install the dependencies and run from this directory, supplying the separate source workbook:

```sh
python -m pip install -r requirements.txt
python cluster_survival.py --workbook ../Supplementary_Data.xlsx --output-dir results
```

The analysis takes a few seconds on a laptop. It writes the 144 treatment-by-parameter means with source-cell addresses, the pairwise Euclidean distances, Ward merges and heights, a results record with the source hash and software versions, and a dendrogram in PNG, PDF and SVG formats. The workbook is read without modification. The numerical reference output is in `reference_results/`.

## Data and analysis

The source is `figure5_survival_raw` in `Supplementary_Data.xlsx`. It contains the survival counts from the Figure 5 simulations. This analysis uses the four plotted life cycles: M100, U25/M75, U100 and U50/M50. Each has nine growth effects and four group-selection effects. Growth multipliers span log2 values −1 to 1 in steps of 0.25; group-selection weights have log2 values 0, 1, 2 and 3.

For each life cycle and parameter combination, we average three survival counts, each from 1,000 runs. The endpoint is survival after eight rounds of growth and transfer. The resulting 36 mean counts form one feature vector per life cycle. Features are equally weighted and are not standardized, centered or transformed. We cluster these four vectors using Euclidean distance and Ward's minimum-variance linkage (`scipy.cluster.hierarchy.linkage`, method `ward`). The square pairwise-distance matrix is an output for inspection; it is not passed as an observation matrix to the clustering function.

Ward linkage gives these merges:

| Clusters joined | Ward height |
| --- | ---: |
| U100 and U50/M50 | 384.129752 |
| U25/M75 and the U100/U50/M50 cluster | 641.425525 |
| M100 and the other three life cycles | 1840.906862 |

Heights are on the mean-survival-count scale, not the probability scale. Dividing every feature by 1,000 preserves the merges and divides all heights by 1,000. This is a descriptive comparison of four modeled response surfaces. The batches and parameter combinations are not additional biological replicates, and the tree does not supply a test of statistical significance.
